# De Novo Design of κ-Opioid Receptor Antagonists Using a Generative Deep Learning Framework

**DOI:** 10.1101/2023.04.25.537995

**Authors:** Leslie Salas-Estrada, Davide Provasi, Xing Qui, H. Ümit Kaniskan, Xi-Ping Huang, Jeffrey F. DiBerto, João Marcelo Lamim Ribeiro, Jian Jin, Bryan L. Roth, Marta Filizola

**Author notes:** Corresponding author: Marta Filizola, PhD, Department of Pharmacological Sciences, Icahn School of Medicine at Mount Sinai, One Gustave L. Levy Place, Box 1677, New York, NY 10029, USA.

## Abstract

Likely effective pharmacological interventions for the treatment of opioid addiction include attempts to attenuate brain reward deficits during periods of abstinence. Pharmacological blockade of the κ-opioid receptor (KOR) has been shown to abolish brain reward deficits in rodents during withdrawal, as well as to reduce the escalation of opioid use in rats with extended access to opioids. Although KOR antagonists represent promising candidates for the treatment of opioid addiction, very few potent selective KOR antagonists are known to date and most of them exhibit significant safety concerns. Here, we used a generative deep learning framework for the *de novo* design of chemotypes with putative KOR antagonistic activity. Molecules generated by models trained with this framework were prioritized for chemical synthesis based on their predicted optimal interactions with the receptor. Our models and proposed training protocol were experimentally validated by binding and functional assays.

## INTRODUCTION

Opioids are widely prescribed medications for the treatment of chronic or severe pain. However, repeated exposure to these drugs can lead to neurological adaptations in the brain that result in addiction.^1, 2^ This chronic, relapsing brain disease, also known as opioid use disorder (OUD), currently affects sixteen million individuals worldwide.^2^ It is characterized by compulsive drug seeking and use despite adverse effects, including respiratory depression, which has recently resulted in approximately 1,000 deaths every week in the United States alone, according to recent estimates.^3^ Addiction to these drugs has led to a global opioid overdose epidemic that has been exacerbated in recent years due to the COVID-19 pandemic.^4-6^ Currently, this ongoing crisis is being addressed from different angles, including prevention of exposure and addiction, support for the discovery of new analgesics, improvement of clinical guidelines for pain management, and the development of treatments for OUD.^7-9^

Suggested priority mechanisms for the development of therapeutics to treat OUDs^10^ include pharmacological blockade of the activity of the LJ-opioid receptor (KOR), a member of the G Protein-Coupled Receptor (GPCR) superfamily. This approach has shown promise in reducing withdrawal symptoms and modulating other key elements linked to addiction such as reward and stress responses.^11-19^ However, the set of known KOR antagonists is limited, and most of them present limitations, including persistent pharmacodynamics, delayed onset of effects, poor brain penetration, and unwanted side effects.^11, 20-24^ Thus, there is a need to broaden the availability of novel KOR antagonists to improve our understanding of the receptor’s function and to develop safer and more effective treatments for opioid addiction.

Recent applications of artificial intelligence (AI)-based tools have demonstrated significant potential in reducing the resources required for traditional drug discovery pipelines, which have estimated cycle times and costs that can range from 10-20 years and 0.8-2.3 billion US dollars, respectively.^25, 26^ Specifically, generative deep learning models have been successfully implemented to solve molecular design and drug development problems.^27-39^ Using architectures composed of neural networks with multiple layers, these models can learn to approximate the probability distribution of the samples used to train them and then to create new samples (not present in the input dataset) from this learned distribution.

In this study, we used a modified version of the Generative Tensorial Reinforcement Learning (GENTRL) architecture^40^ to design KOR antagonists *de novo*. The architecture comprises a conditional variational autoencoder (cVAE) that uses a recurrent neural network (RNN) encoder and a convolutional neural network (CNN) decoder. Recently, this architecture was shown to facilitate the rapid discovery of inhibitors of the discoidin domain receptor 1 from design to *in vivo* verification^40^ by utilizing a two-step protocol consisting of pre-training followed by reinforcement learning (RL) cycles. Here, we first conducted pre-training on a dataset of millions of drug-like compounds in Simplified Molecular Input Line Entry System (SMILES) format to learn an approximation of the distribution of the high-dimensional chemical space, and then on a smaller, specialized dataset that included known KOR inhibitors from the ChEMBL 30 database,^41^ in an attempt to fine-tune the model. The high-dimensional chemical space was mapped by the cVAE into a low-dimensional latent space that encoded the relationships between SMILES and selected molecular properties for use in the generative model to sample compounds within the same learned distribution but beyond the known chemical space. An established reinforcement learning algorithm was then used to bias compound generation towards new chemotypes that were predicted to inhibit KOR’s activity. Five molecules were prioritized for experimental testing based on their high ligand-receptor structural interaction fingerprints (SIFt) and shape similarity to the JDTic crystallographic binding pose at KOR.^42^ These molecules were synthesized and experimentally tested using binding and functional assays. All five compounds were able to bind to the receptor, and the two with the highest binding affinity were functionally evaluated and found to exhibit antagonism at KOR.

## MATERIALS AND METHODS

### Generative Deep Learning Architecture

The conditional variational autoencoder within the GENTRL’s open framework^40^ was used as the basis for our generative modeling strategy. The encoder consisted of a 2-layer RNN with gated recurrent units of hidden size 256, while the decoder consisted of a 128-channel dilated CNN with 7 stacked layers. A 50-dimensional latent space was parameterized using tensor decomposition with mixtures of 20 Gaussian components at each dimension and tensor-trains with a core size of 30. The architecture was modified to correctly encode and decode all entries in the training datasets by extending the vocabulary to include all the necessary characters and increasing the maximum allowed length of individual entries (see below).

### Pre-Training Datasets, Properties, and Evaluation

We used two different datasets to pre-train the generative model. Dataset #1 consisted of a curated list of 4.6 million compounds. To obtain this curated dataset, we started with the 6.2 million two-dimensional (2D), clean, in-stock, drug-like compounds from the ZINC 20 database^43^ (as of 03/08/2021) and removed molecules that had atoms besides C, N, S, O, F, Cl, Br, and H, cycles with more than 8 atoms, more than 7 rotatable bonds, and potentially toxic and/or reactive moieties based on medicinal chemistry filters (MCFs) and Pan Assay Interference Compounds (PAINS) filters.^44-46^ Next, we split the curated set into a pre-training dataset (80%), a test set (10%), and a scaffold test set (10%). The latter was composed of a set of molecules with Bemis-Murcko scaffolds not present in either the pre-training dataset or the test set and was used to assess the generative model’s ability to produce molecules with novel scaffolds.^47, 48^ Dataset #2 consisted of 2,056 compounds from the ChEMBL 30 database^41^ (as of 01/27/2022) with reported half-maximum inhibitory concentration (IC_50_) at human KOR (single-protein target ChEMBL ID 237). The dataset was divided into two categories: “positive” KOR inhibitors (IC_50_ < 1 µM) and “negative” KOR inhibitors (IC_50_ ≥ 1µM), consisting of 445 and 1,611 molecules, respectively. Both pre-training datasets were saved in the SMILES format.

Five molecular properties were calculated for all molecules in the pre-training datasets: 1) Topological polar surface area (TPSA), estimated using the RDKit’s TPSA descriptor based on contributions from N, O, S, and P atoms;^49, 50^ 2) Lipophilicity as per Wildman-Crippen’s log *P* (Wlog *P*), calculated using RDKit’s MolLogP descriptor with an atom-based approach;^50, 51^ 3) Solubility as per log *S*, estimated using Delaney’s Estimated SOLubility (ESOL) method;^52^ 4) A binary flag indicating whether the molecule passed (1) or failed (0) MCFs; and 5) A binary flag indicating whether the molecule was a KOR inhibitor (1) or a non-inhibitor (0), based on its reported IC_50_ at KOR. This property was only available for molecules in Dataset #2 and their corresponding entries in Dataset #1. For molecules without IC_50_ records, the corresponding property was left empty to indicate it as missing.

The model was initially pre-trained on Dataset #1 for up to 25 epochs using an Adam optimizer with a learning rate of 10^-4^. Each epoch represents one pass over the complete dataset in training batches of 50 SMILES.^53^ The quality of the model and its ability to approximate the distribution learned from Dataset #1 was assessed using established metrics^48^ and 30,000 valid SMILES sampled after 0, 2, 4, 6, 8, 10, 15, 20, and 25 epochs of pre-training. These metrics included validity, uniqueness, internal diversity, and passing chemical filters. Validity was assessed by verifying successful/failed conversion to RDKit’s and OpenEye’s molecule objects.^50, 54^ Uniqueness was defined as the fraction of valid SMILES that were not duplicated in a generated set of molecules.^48^ Internal diversity was measured as 1 minus the average chemical similarity as per Tanimoto coefficient (Tc) within the generated batch, using Morgan fingerprints.^48^ The same chemical filters used during the construction of pre-training Dataset #1 were used to evaluate the fraction of molecules passing chemical filters. The generated molecules were also compared to the reference datasets (test set and scaffolds test set) using established metrics to assess the quality of generative models.^48^ These metrics included BRICS fragments similarity,^55^ Bemis-Murcko scaffolds^47^ similarity, molecular properties (e.g., molecular weight, lipophilicity, synthetic accessibility, and quantitative estimate of drug-likeness), as well as the Fréchet ChemNet Distance (FCD)^56^ and average Tanimoto similarity of Morgan fingerprints to the nearest neighbor in the reference sets.

In the second step of pre-training, the model underwent 300,000 updates using an Adam optimizer with a learning rate of 10^-4^. An update consists of a pass over one batch of 200 molecules, which included 100 molecules from Dataset #1, 50 molecules from the “positive” set in Dataset #2, and 50 molecules from the “negative” set in Dataset #2.

### Reinforcement Learning

Reinforcement learning was performed using the REINFORCE algorithm^57^ within the GENTRL’s reinforcement learning framework. Training was carried out for 9,000 iterations, using an Adam optimizer with a learning rate of 10^-5^ and a batch size of 200. An iteration consists of one batch of 200 SMILES generated and scored by the model. The reward function was computed as the product of six reward components, calculated as follows:

1) *Pharmacophoric similarity (Pharm).* The OpenEye Python Toolkits (version 2021.2.0)^54^ were used to generate 3D conformers of SMILES, following the protocol outlined in ref. ^58^. Briefly, the most probable tautomeric form and protonation state at pH 7.4 were determined using the Quacpac Toolkit (version 2.1.3.0),^59^ while 3D structure generation was carried out with the Omega Toolkit (version 4.1.2.0),^60^ which performed stereoisomer enumeration for up to 3 stereocenters and a maximum of 200 conformers. To assess similarity to JDTic’s binding mode at KOR (PDB ID 4DJH),^42^ two methods were used: a) the Ultrafast Shape Recognition with CREDO Atom Types score^61^ (USRCAT score) or b) the Shaper2^62^ alignment score to cavity-derived pharmacophoric features determined with the IChem toolkit^63^ (IChem score). To calculate USRCAT scores, the OpenEye-generated 3D conformers were converted into RDKit molecule objects using the Python module available in ref. ^64^ and then ranked according to their URSCAT scores. To calculate IChem scores, chain B of the human inactive JDTic-bound KOR structure (PDB ID 4DJH)^42^ was prepared using the default protocol of the Protein Preparation Wizard in the Schrödinger 2020-4 suite.^65^ The protocol consisted of: (i) back mutating residue L135 to isoleucine to match the wild-type sequence; (ii) adding missing side chains and hydrogen atoms; (iii) capping the N-and C-term of the chain at residues S55 and P347, respectively; (iv) assigning correct bond orders and protonation states for the ligand and the side chains at pH 7.4; and (iv) conducting a short, restrained energy minimization *in vacuo* using the OPLS3e force field.^66^ Then, mol2 files of the prepared KOR and JDTic structures were generated using ChimeraX^67, 68^ and the IChem’s Volsite tool was used in the ligand-restricted mode to derive pharmacophoric features from the cavity around the ligand, as described in ref. ^62^. The overlap of conformers to the cavity-based pharmacophore was assessed with the Shaper2 tool using default parameters. The Pharm reward component was defined as either the best USRCAT or IChem score among the generated conformers.
2) *Medicinal chemistry filters (MCFs).* MCFs were evaluated using substructure matching against a library of SMARTS patterns from the Molecular Sets (MOSES) platform.^48^ The MCF reward component was defined with a binary flag that indicated whether a molecule passed (1) or (0) failed the filters.
3) *Blood-brain barrier permeability (BBB).* BBB was estimated using the Brain Or IntestinaL EstimateD permeation (BOILED-Egg) method, which is a graphical model that predicts brain access based on a molecule’s Wlog *P* and TPSA.^49, 69^ The BBB reward component was defined with a binary flag that indicated whether a molecule was predicted to be BBB penetrant (1) or not (0).
4) *Solubility (log S).* Log S was estimated using the ESOL method.^52^ This method is based on a linear regression model derived from ∼2,900 solubility measurements that include parameters such as log *P*_octanol_, molecular weight, proportion of heavy atoms in aromatic systems, and number of rotatable bonds. The solubility reward component was defined with a binary flag that indicated whether the predicted log *S* was between -5 and -1^70^ (1) or not (0).
5) *Ease of synthesis (SA)*. SA was evaluated using Ertl and Schuffenhauer’s synthetic accessibility score (SAS),^71^ which ranks molecular complexity based on a library of substructures on a scale of 1 (easiest) to 10 (hardest), where SAS > 6 is considered difficult to synthesize.^49^ The SA reward component was defined as the normalized value (10 - SAS)/9, where a score of 0 indicates the hardest molecules to synthesize and 1 indicates the easiest.^72^
6) *Novelty against known KOR inhibitors (novelty).* Novelty was assessed based on the maximum Tanimoto coefficient (Tc_max_) against the Morgan topological 2D fingerprints of molecules in the “positive” subset of Dataset #2, which consisted of 445 compounds with IC_50_ < 1 µM at KOR. The novelty reward component was defined as 1 – Tc_max_, where a score of 0 indicates that the molecule is not novel and a score of 1 indicates novel molecules.

A reward of 0 was assigned to invalid molecules, as determined by the failed conversion of SMILES to RDKit and/or OpenEye molecule objects,^50, 54^ or to molecules for which the generation of 3D conformers was unsuccessful.

### Filtering

The generated molecules were scored using the six reward components defined in the reinforcement learning stage and selected for docking if they met the following criteria: a USRCAT score > 0.25 or an IChem score > 0.4, passing MCFs, likely BBB permeability, -5 ≤ log *S* ≤ -1, SAS < 6, and Tc_max_ < 0.5.

### Molecular Docking

The molecules were prepared for docking with the LigPrep utility included in the Schrödinger 2020-4 suite.^65^ The ionization state and tautomeric forms determined by the pharmacophoric similarity reward component were preserved during ligand preparation. The human inactive JDTic-bound KOR structure (PDB ID 4DJH)^42^ was prepared for docking using the default protocol of the Protein Preparation Wizard in the Schrödinger 2020-4 suite,^65^ as described above. A cubic docking grid with a edge of 26 Å was generated on the center of mass of JDTic using Schrodinger’s Receptor Grid Generation utility.^65^ The molecules were docked using Glide with the Extra Precision (XP) scoring function,^65^ and JDTic was docked using the same procedure as a control.

### Structural Interaction Fingerprints (SIFts)

An in-house python script was used to calculate the following types of protein-ligand interactions between the docked molecules and KOR residues: 1) carbon-carbon interactions within 4.5 Å (Apolar), 2) face-to-face aromatic interactions within 4 Å (Aro_F2F), 3) edge-to-face aromatic interactions within 4 Å (Aro_E2F), 4) hydrogen-bonds with a protein residue as the donor (Hbond_ProD), 5) hydrogen-bonds with a protein residue as the acceptor (Hbond_ProA), 6) electrostatic interactions (within 4 Å) with a protein residue positively charged (Elec_ProP), 7) electrostatic interactions (within 4 Å) with a protein residue negatively charged (Elec_ProN). The similarity of the poses of the docked molecules to JDTic’s pose in the JDTic-bound crystal structure (PDB ID 4DJH)^42^ was assessed by Tanimoto coefficients of their respective SIFts using RDKit.^50^

### Chemical Synthesis

Common materials or reagents were purchased from commercial sources and used without further purification. High-performance liquid chromatography (HPLC) spectra were acquired using an Agilent 1200 Series system with DAD detector for all the intermediates and final products below. Chromatography was performed on a 2.1×150 mm Zorbax 300SB-C18 5 µm column with water containing 0.1% formic acid as solvent A and acetonitrile containing 0.1% formic acid as solvent B at a flow rate of 0.4 ml/min. The gradient program was as follows: 1% B (0−1 min), 1−99% B (1−4 min), and 99% B (4−8 min). High-resolution mass spectra (HRMS) data were acquired in positive ion mode using an Agilent G1969A API-TOF with an electrospray ionization (ESI) source. Ultra-performance liquid chromatography (UPLC) spectra for compounds were acquired using a Waters Acquity I-Class UPLC system with a photo diode array (PDA) detector. Chromatography was performed on a 2.1 Å∼ 30 mm Acquity UPLC BEH C18 1.7 μm column with water containing 3% acetonitrile, 0.1% formic acid as solvent A and acetonitrile containing 0.1% formic acid as solvent B at a flow rate of 0.8 mL/min. The gradient program was as follows: 1−99% B (1−1.5 min), and 99−1% B (1.5−2.5 min). Preparative HPLC was performed on Agilent Prep 1200 series with UV detector set to 220 or 254 nm. Samples were injected onto a Phenomenex Luna 250 x 30 mm, 5 µm, C_18_ column at room temperature. The flow rate was 40 ml/min. A linear gradient was used with 10% of acetonitrile in H_2_O (with 0.1 % TFA) (B) to 100% of acetonitrile (A). Nuclear Magnetic Resonance (NMR) spectra were acquired on a Bruker DRX-400 spectrometer with 400 MHz for proton (^1^H NMR) or a Bruker Avance-III 800 MHz spectrometer with 800 MHz for proton (^1^H NMR); chemical shifts are reported in (δ). HPLC was used to establish the purity of target compounds. All final compounds had > 96% purity using the HPLC methods described above. The chemical synthesis of compounds (S)-N-((1-(3,4-difluorobenzyl)piperidin-3-yl)methyl)-4,6-dimethylpicolinamide (FJRL-14), 2-((2-chloro-4-methylbenzyl)(methyl)amino)-1-(4-(4-chlorobenzyl)-1,4-diazepan-1-yl)ethan-1-one (FJRL-20), (S)-N-((1-(3,4-difluorobenzyl)piperidin-3-yl)methyl)-4,6-dimethylpicolinamide (FJRL-29), 1-((6-methylpyridin-2-yl)methyl)-N-(2-(pyridin-3-yl)ethyl)azepane-4-carboxamide (FJRL-36), and N-((1-(4-isopropylbenzyl)piperidin-4-yl)methyl)-4-methylbenzamide (FJRL-91) is described in detail in Supporting Information.

### Competition Binding Assays

Radioligand binding assays were carried out using membranes prepared from HEK293 cells transiently transfected with human KOR. Samples were mixed with KOR membranes and ^3^H-U69593 (∼1 nM) in 96-well plates. After incubation of 1 hr at room temperature, the radioligand bound to KOR membranes was filtered and separated from the free radioligand. Radioactivity was measured with a MicroBeta counter. Results were compared to reference Salvinorin A (Sal A) (0 – 100%) and fitted to the built-in One-site function for competitive binding assays in GraphPad Prism (v9.5).^73^

### Cell-Based Functional Assays

G_i_ GloSensor cAMP assays were carried out in transiently transfected HEK293 T cells. Overnight transfected cells (KOR + GloSensor) were plated in poly-L-Lys coated 384-well white plates in DMEM supplemented with 1% dialyzed FBS for 6 hours. After the medium was removed, cells were stimulated with samples premixed with luciferase substrate for 10 min at room temperature, followed by addition of isoproterenol (100 nM) to activate endogenous β_2_-adrenergic receptor-stimulated G_s_ signaling. The plate was then read after 20 min by a luminescence reader. For antagonist activity, 10 nM Sal A was added at 10 min after samples for additional 10 min incubation before isoproterenol. Results (Relative Luminescence Units) were normalized to basal (1.0 fold of basal) and were analyzed with the built-in four-parameter concentration-response function in GraphPad Prism.^73^ Reduction in luminescence is proportional to G_i_ activation.

## RESULTS

### Distribution Learning and *De Novo* Compound Generation

Figure 1 shows a schematic representation of the generative deep learning strategy that we used to design KOR ligands with antagonistic activity, leveraging the structural and physicochemical information of the receptor, its ligands, and large chemical libraries of drug-like molecules.

**Figure 1.**
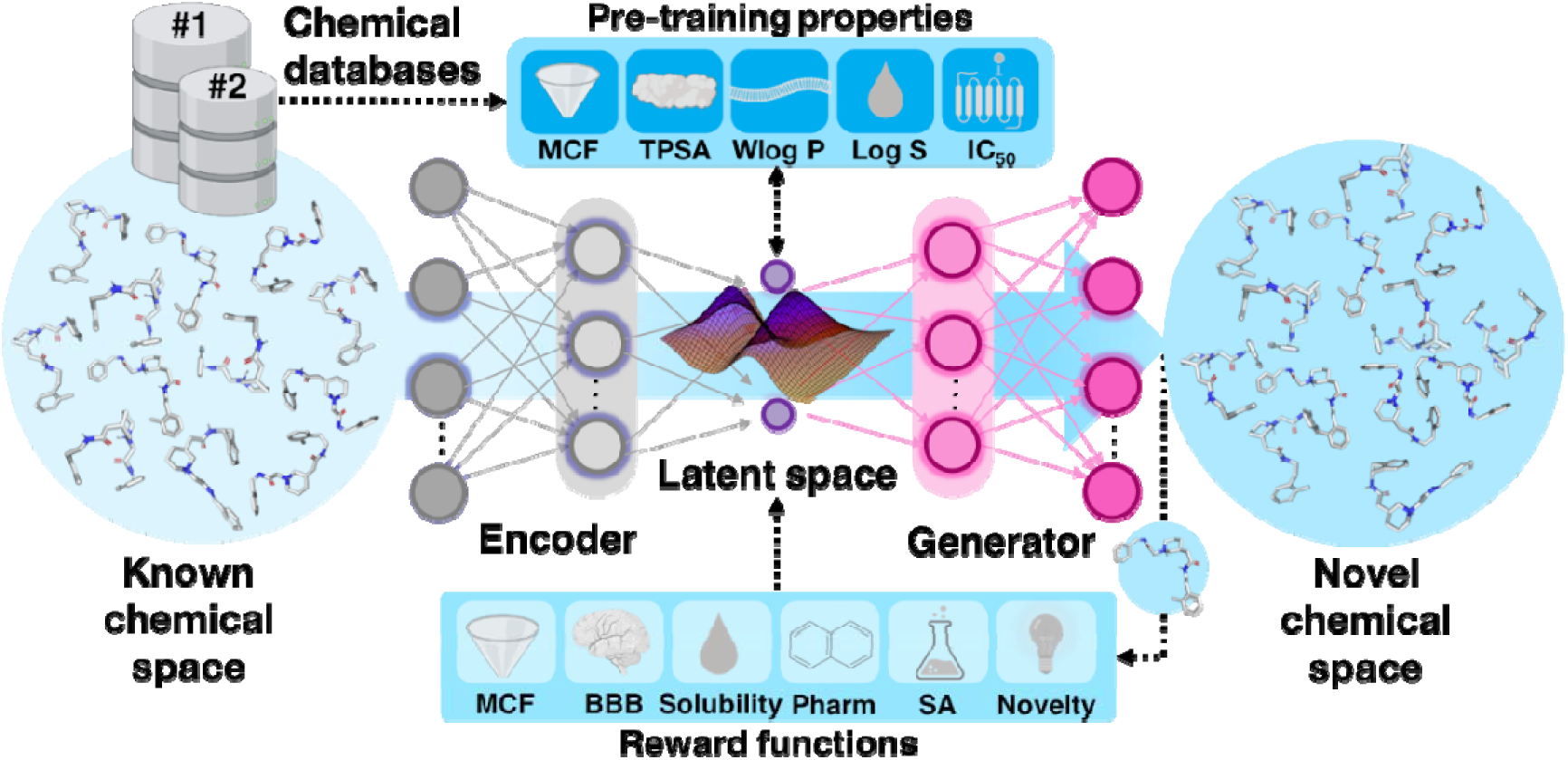
Scheme of the generative deep learning strategy used for the *de novo* design of KOR antagonists. MCF: medicinal chemistry filters; Wlog *P*: Wildman-Crippen’s log *P*; Log *S*: logarithm of solubility; BBB: blood-brain barrier; Pharm: pharmacophore similarity; SA: ease of synthesis.

Specifically, we used a modified version of the GENTRL architecture^40^ and pre-trained a model using data derived from 4.6 million molecules from the ZINC 20 database^43^ (Dataset #1). These molecules were characterized by molecular properties such as MCFs,^48^ TPSA, Wlog *P*, log *S,*^52^, ^70^ and reported KOR inhibition (IC_50_) (see Methods for further details). We evaluated the performance of this model by conducting pre-training for an increasing number of epochs. Each epoch involved the model passing over the entire Dataset #1 once. We assessed the model’s ability to generate chemically valid, novel, and diverse drug-like compounds and to approximate the distribution learned from Dataset #1 using established metrics^48^ (see Methods for details). Figure S1 illustrates the model’s performance in generating 30,000 valid SMILES as the number of pre-training epochs increases. The highest validity values were achieved after 20 epochs, and we selected the resulting model for fine-tuning via a second round of pre-training using Dataset #2. This dataset consisted of 300,000 batches, each containing 100 molecules from Dataset #1, 50 KOR inhibitors, and 50 KOR non-inhibitors (see Methods for details). The performance of the model in producing 30,000 valid SMILES after the second round of pre-training is also depicted in Figure S1, whereas Tables S1-S2 provide estimates of model’s performance on reference datasets after pre-training on Dataset #1 or #2, respectively. The parameters obtained from the model after 20 epochs were saved and used as the starting point for a reinforcement learning step.

### Reinforcement Learning for *De Novo* Generation of KOR Antagonists

Two separate models were trained using a multi-component reward function computed as the product of the reward components described in the Methods section. These components included an estimate of ligand-based or cavity-based pharmacophoric similarity to the ligand binding mode in the JDTic-bound KOR structure,^42^ expressed as a USRCAT score or IChem score, respectively. The performance of the two models in producing 30,000 valid molecules was monitored during reinforcement learning by a set of indicators, including the fraction of molecules with a combined reward greater than 0.1 (Figure 2), as well as the validity of chemical structures, their internal diversity, uniqueness, and passing of chemistry filters (Figure S2). The training improved learning in both models up to 9,000 reinforcement learning iterations, but incorporating cavity-based information produced a larger fraction of valid molecules.

**Figure 2.**
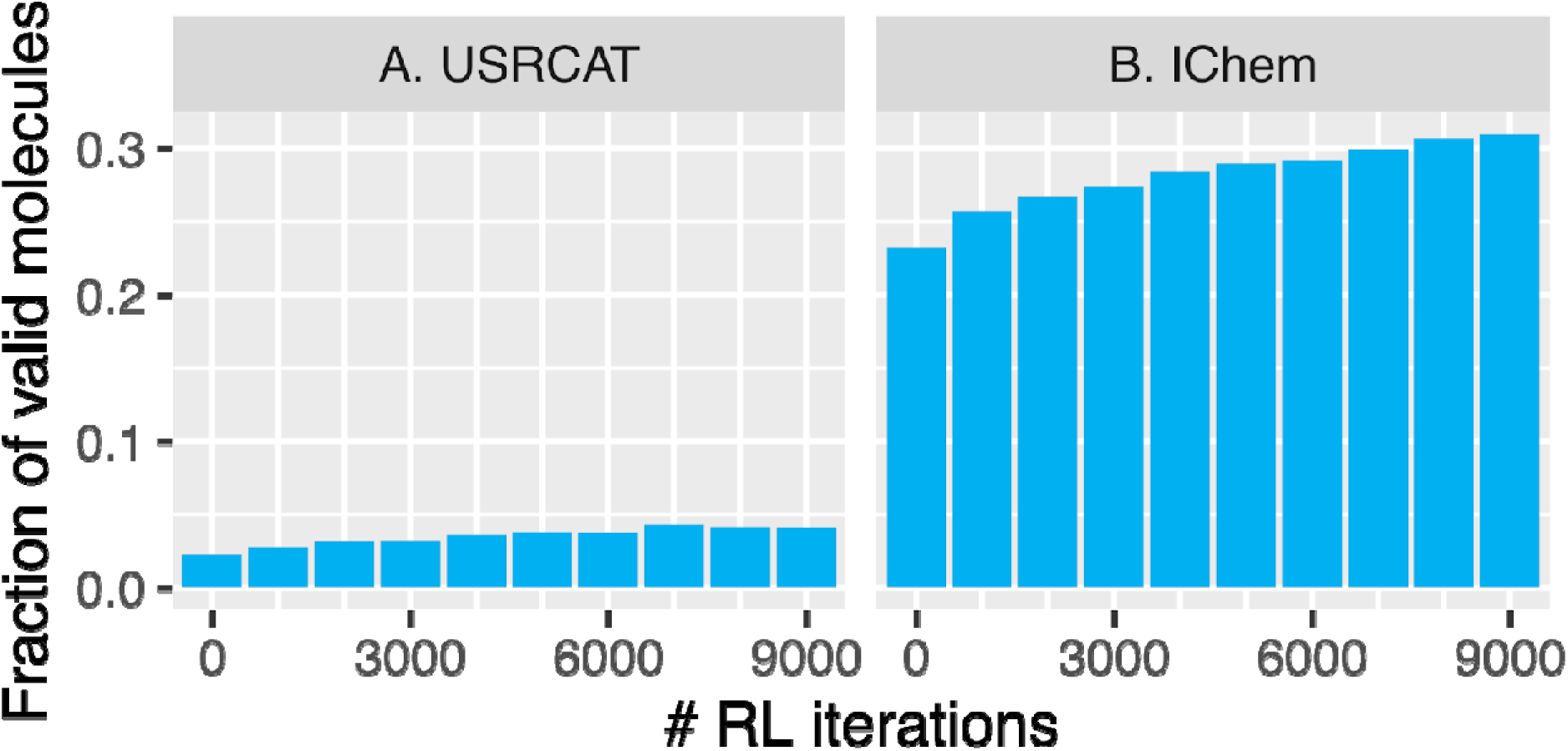
Performance of the generative deep learning models in producing molecules highly scored by a multi-component reward function. A) Model trained using ligand-base pharmacophoric similarity assessed by USRCAT scores. B) Model trained using cavity-base pharmacophoric similarity assessed by IChem scores. Each reinforcement learning iteration consists of the model generating and scoring 200 compounds in SMILES format.

### Compound Prioritization for Chemical Synthesis and Experimental Testing

Figure 3 illustrates the strategy used to prioritize molecules for chemical synthesis and experimental testing. Firstly, the two trained models were used to generate sets of 500,000 valid SMILES each after 9,000 reinforcement learning iterations, resulting in a total of 1 million SMILES. Compounds that met the desired reward values for ligand-based or cavity-based pharmacophore similarity (USRCAT > 0.25 or IChem > 0.4, respectively), novelty (Tc_max_ < 0.5), ease of synthesis (SAS < 6), solubility (-5 ≤ log *S* ≤ -1), and likelihood to penetrate the BBB while passing MCFs were selected for molecular docking. In total, 2,545 compounds satisfied these criteria, with 1,116 and 934 molecules derived from the model using USRCAT and IChem scores, respectively. The best 3D conformers of these molecules, as determined by the pharmacophoric similarity reward component, were prepared for docking at KOR (PDB ID 4DJH)^42^ in the same ionization state and tautomeric form. The docked compounds using Glide XP^65^ and JDTic as a docking control were ranked by their similarity to JDTic’s interactions with KOR. Table S3 shows the 2D structures of the 169 molecules exhibiting the highest similarity to JDTic’s interactions with KOR, as assessed by SIFt Tc values larger than 0.75. Notably, more molecules generated by the model using IChem scores had ligand-receptor interactions closer to JDTic’s than those generated by the model using USRCAT scores (116 vs. 53 compounds with SIFt Tc ≥ 0.75).

**Figure 3.**
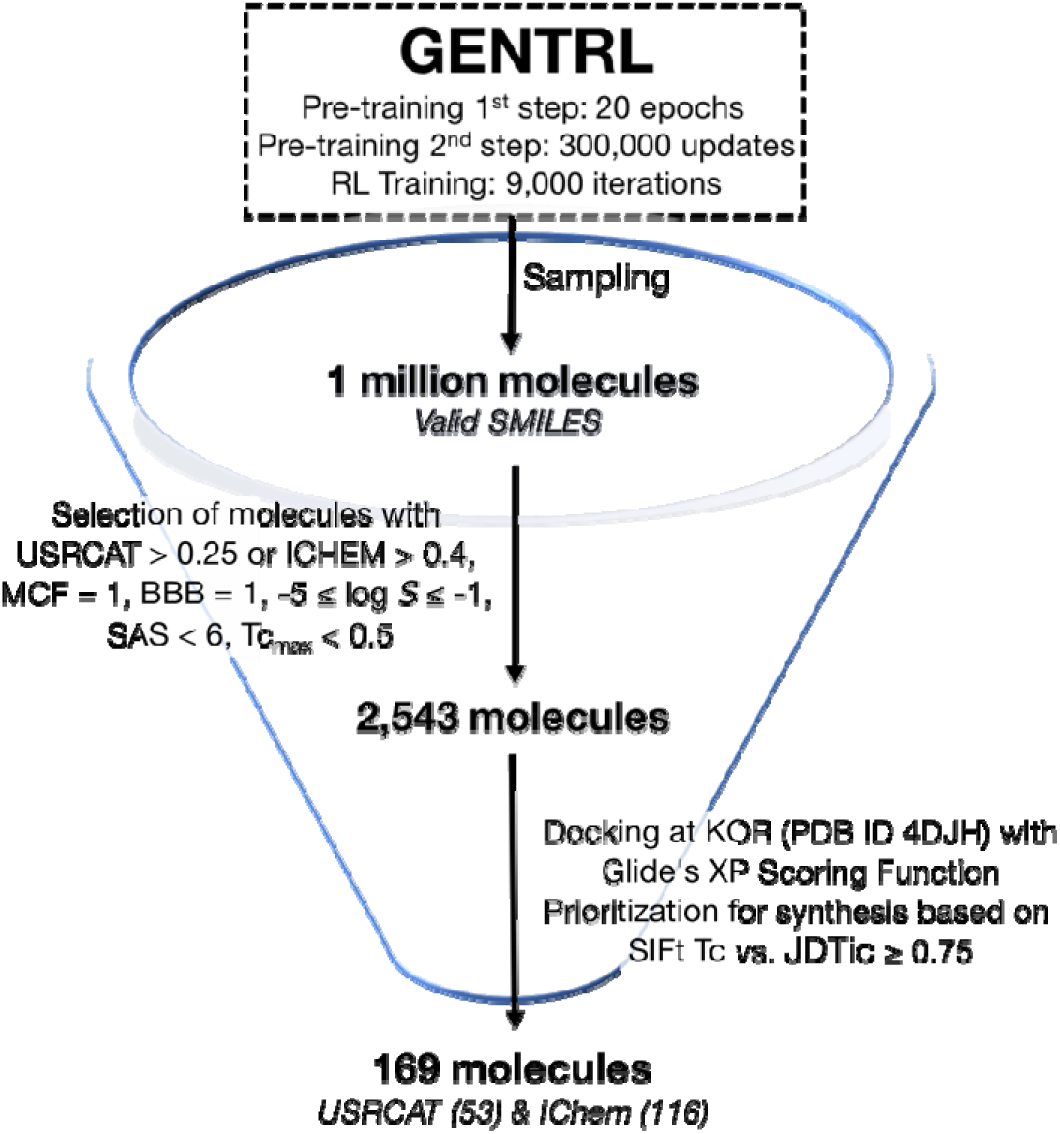
Prioritization pipeline for chemical synthesis and experimental testing of compounds generated by models trained with reinforcement learning. Two sets of 500,00 compounds each were generated from models using USRCAT and IChem scores.

Five top-scoring docked compounds (FJRL-14, FJRL-20, FJRL-29, FJRL-36, and FJRL-91) based on SIFt Tc values were selected for chemical synthesis and experimental validation based on the readily availability of chemical materials. Details of the chemical synthesis for these compounds are reported in the Materials and Methods section and Supporting Information. As expected based on their similarity in binding mode with JDTic, all five compounds were predicted to have salt-bridge or hydrogen-bond interactions with KOR residue D138^3.32^ in transmembrane (TM) 3 (Figure S3). Superscript numbers here and throughout refer to the Ballesteros-Weinstein’s generic numbering scheme for GPCRs.^74, 75^ Additionally, all compounds shared hydrophobic interactions with residues in TM2 (V108^2.53^), TM3 (V134^3.28^, I135^3.29^, D138^3.32^, Y139^3.33^, M142^3.36^), TM6 (W287^6.48^, I290^6.51^, I294^6.55^), and TM7 (I316^7.39^ and Y320^7.43^) (Figure S4).

### Experimental Validation of Synthesized Compounds

Competition binding assays were performed for compounds FJRL-14, FJRL-20, FJRL-29, FJRL-36, and FJRL-91 according to the protocols described in the Materials and Methods section. All five tested compounds were found to bind to the receptor at high concentrations with K_i_ values of 6.46 μM (pK_i_ of 5.19 ± 0.06) and 7.59 μM (pKi of 5.12 ± 0.06) for FJRL-14 and FJRL-20, and more than 10 µM for FJRL-29, FJRL-36, and FJRL-91, respectively, compared to JDTic and Salvinorin A, which were used as positive controls (pK_i_ = 8.45 ± 0.10, K_i_ = 3.55 nM, and pK_i_ = 8.30 ± 0.13, K_i_ = 5.01 nM, respectively). Figure 4 shows the two compounds with the highest binding affinity for KOR, specifically, FJRL-14 and FJRL-20, docked at the KOR binding pocket (Figures 4A and 4D, respectively), as well as their results from competition binding assays using KOR membranes and 1 nM ^3^H-U69593 (Figures 4B and 4E, respectively), and Sal A-mediated G_i_ inhibition of cAMP production as assessed in transiently transfected HEK293 T cells (Figure 4C and 4F, respectively). FJRL-14 and FJRL-20 showed no agonist activity (results not shown) but weak antagonist activity against Sal A-mediated G_i_ activation. Details of the experimental protocols are provided in the Materials and Methods section.

**Figure 4.**
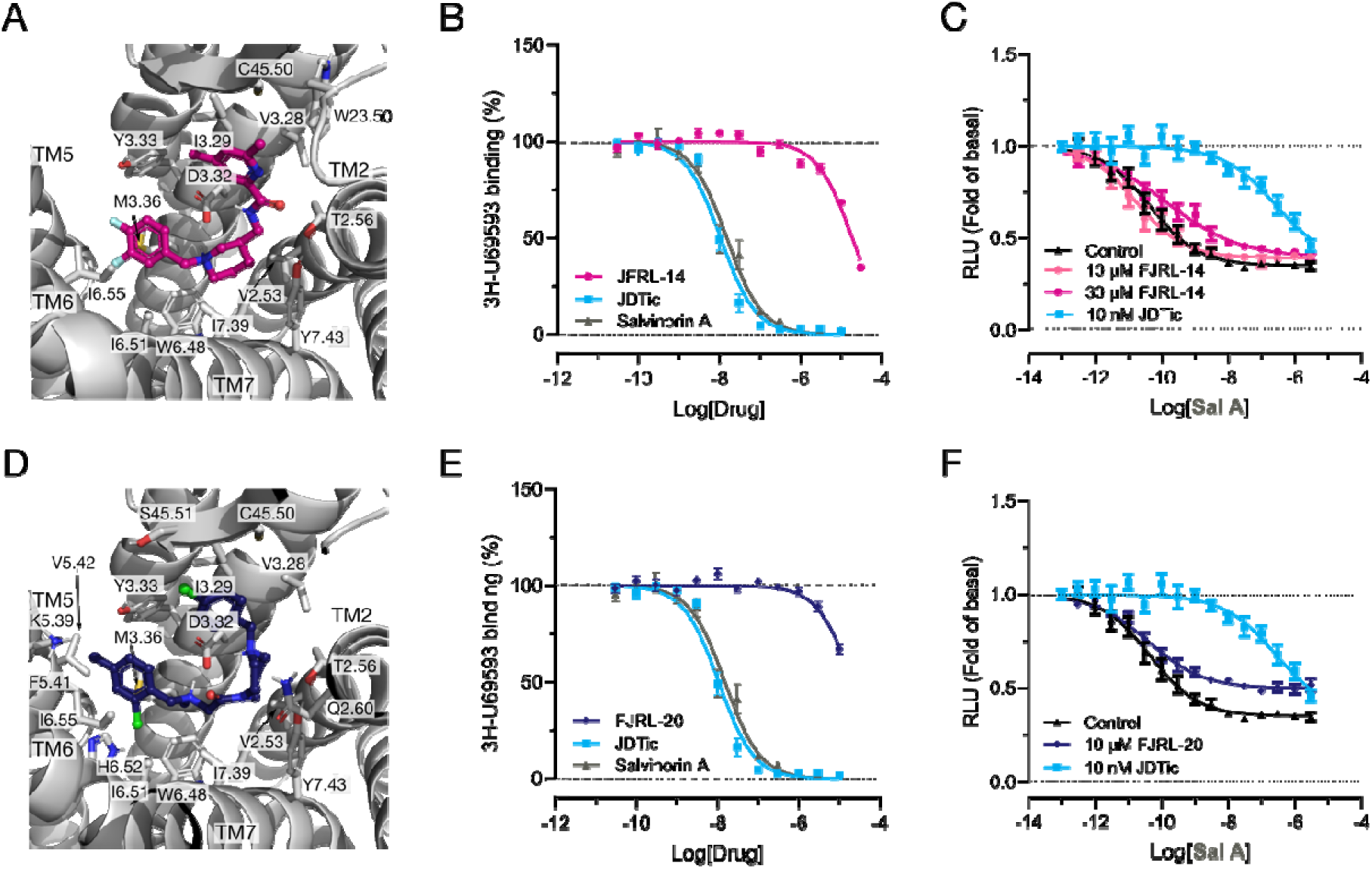
*De novo*-generated KOR antagonists FJRL-14 and FJRL-20. (A, D) Docked compounds at the JDTic-KOR’s crystal structure (PDB ID 4DJH) with interacting residues as per structural interaction fingerprints shown as sticks. (B, E) Competition binding assays using KOR membranes and 1 nM ^3^H-U69593 for FJRL-14 and FJRL-20, respectively. (C, F) KOR-mediated Gi inhibition of cAMP production as assessed in transiently transfected HEK293 T cells using GloSensor cAMP assay (Promega, USA) for FJRL-14 and FJRL-20, respectively.

As also shown in Figure S3, the predicted binding mode of FJRL-14 and FJRL-20 at KOR is very similar to JDTic’s, with the ligands sharing most ligand-receptor interactions (Figure S4). However, despite this high level of similarity, at least one of the two compounds is missing interactions with KOR that are present for JDTic (e.g., Q115^2.60^, G319^7.42^) and are not replaced by an equivalent number of new interactions. This reduced number of interactions between the *de novo* compounds and the receptor is likely responsible for the observed lower binding affinities as per radioligand binding assays and weaker antagonistic activities as determined by Schild analysis of competitive antagonism. Confirmation of the novelty of FJRL-14 and FJRL-20, in comparison to the reported 337 opioid ligands in the IUPHAR/BPS Guide to Pharmacology database,^76^ was achieved by calculating Tc_max_ values below 0.5 and examining the closest 2D structures. A chemical similarity search between FJRL-14 and FJRL-20 with the entire ChEMBL 30 database^41^ (∼2.2. million compounds) resulted in Tc_max_ values of 0.67 and 0.70, respectively, providing further evidence of their chemical novelty.

## DISCUSSION

In recent years, AI-based tools have shown great potential for enhancing drug research and development, particularly through the use of generative deep learning architectures. However, the validation of AI-assisted novel molecule identification remains a crucial challenge, especially for central nervous system diseases, often due to a shortage of high-quality data to train reproducible ML models, difficulties in interpreting data derived from ML models, and the challenging need to integrate data from various sources. Despite these limitations, our work demonstrates that deep generative models offer a promising solution to accelerate the early lead discovery process for members of the GPCR superfamily.

In our study, we used a generative modeling architecture that had been experimentally shown to generate potent leads in just 21 days from conception to *in vivo* validation. However, we had to modify it to adapt it to our specific problem and to compensate for the lack of accessibility to proprietary information, including the code for the reinforcement learning reward functions. Thus, we utilized publicly available databases of chemical compounds from which to derive drug-like properties, as well as structural and functional information about KOR, to establish and validate a training protocol for generating novel KOR antagonists. Our work demonstrates that this revised protocol and algorithm enables the exploration of uncharted regions of the chemical space and provides a viable alternative to biased chemical library screening.

While the compounds we have been able to synthesize and test so far exhibited weak antagonistic activity, our results confirm that these tools can effectively design new compounds with desired pharmacological profiles *in vitro*. However, further studies are needed to verify whether these molecules also have desirable pharmacological profiles *in vivo*, such as BBB permeability, which we included as a reward function. Therefore, we are currently working on incorporating potency into our generative architecture to enhance the possibility of obtaining potent compounds that can be candidates for animal testing.

One area where we see the potential for further improvement is increasing the yield of valid molecules, although this limitation could be circumvented by increasing the number of molecules being generated. The current low yield of valid molecules is likely due to using a RNN for the encoder, which becomes harder to train with longer input sequences. In addition to using different encoders, testing different physicochemical properties and structural evaluations for training could also improve the yield of valid molecules.

Additionally, we found that using reward functions derived from cavity-based information allowed us to generate more molecules with ligand-receptor interactions closer to JDTic’s than reward functions derived from ligand-based molecular shape features. Future work is required to investigate different hybrid ligand-and structure-based training schemes.

It is worth noting that our protocol can be generalized to other GPCRs with available three-dimensional structures or molecular models by updating the datasets used for training with sets that are relevant to the GPCR under study. As new structural and functional information of GPCRs becomes available, we expect that the performance of AI-driven molecular design models will continue to improve and have further impact in reducing the times and costs associated with drug discovery for the treatment of various conditions, including opioid addiction.

## CONCLUSIONS

While there are challenges associated with applying AI/ML technology to drug discovery, our work highlights the potential benefits of using deep generative models to accelerate the early lead discovery process for GPCRs. Our study demonstrates that AI-driven molecular design can effectively generate novel compounds with desired pharmacological profiles *in vitro*, and we anticipate that continued improvements in AI-driven molecular design models will lead to further advancements in drug discovery for a range of diseases.

## DATA AND MATERIAL AVAILABILITY

GENTRL and MOSES were obtained from their respective GitHub repositories (https://github.com/insilicomedicine/GENTRL and https://github.com/molecularsets/moses). Chemical libraries were downloaded from the ZINC 20 database (https://zinc20.docking.org/) and the ChEMBL 30 database (https://www.ebi.ac.uk/chembl/). PDB files were downloaded from the RCSB Protein Data Bank (https://www.rcsb.org). Glide XP within the Schrödinger 2020-4 suite was used for docking. SIFt analysis was carried out using an in-house Python script which can be made available upon request. PyMOL(TM) 2.5.2 was used for molecular visualization (https://pymol.org/2/). GraphPad Prism 9.5 was used for analysis of binding and functional assays. All data generated or analyzed during this study are available from the corresponding authors upon request.

## SUPPORTING INFORMATION

Details of chemical synthesis of selected compounds; Performance of the deep generative model using reference datasets after pre-training on Dataset #1 (Table S1); Performance of the deep generative model using reference datasets after pre-training on Dataset #2 (Table S2); List of 169 compounds prioritized for chemical synthesis (Table S3); Performance of the deep generative model in producing chemically valid, novel, and diverse molecules during the pre-training steps (Figure S1); Performance of the deep generative model in producing chemically valid, novel, and diverse molecules during the reinforcement learning step (Figure S2); Compounds prioritized for chemical synthesis docked at KOR and compared to JDTic’s binding mode (Figure S3); Protein-ligand interactions of compounds prioritized for chemical synthesis docked at KOR and compared to JDTic’s (Figure S4); Simplified Molecular Input Line Entry System (SMILES) for compounds presented in the manuscript (SMILES.csv).

## ACKNOWLEDGMENTS

This work was funded by National Institutes of Health grant DA045473. Computations were supported in part through the computational resources and staff expertise provided by Scientific Computing at the Icahn School of Medicine at Mount Sinai, the Clinical and Translational Science Awards (CTSA) grant UL1TR004419 from the National Center for Advancing Translational Sciences, and the Office of Research Infrastructure of the National Institutes of Health under award number S10OD026880. The work utilized the NMR Spectrometer Systems at Mount Sinai acquired with funding from the NIH’s Shared Instrumentation Grants 1S10OD025132 and 1S10OD028504. Radioligand binding and GloSensor cAMP assays were performed under the National Institute of Mental Health’s Psychoactive Drug Screening Program (NIMH PDSP), contract HHSN-271-2018-00023-C. The NIMH PDSP is directed by Bryan L Roth, MD, PhD at University of North Carolina at Chapel Hill and Project Officer Jamie Driscoll at NIMH, Bethesda, MD, USA. The content is solely the responsibility of the authors and does not necessarily represent the official views of the National Institutes of Health.

## For Table of Contents Use Only

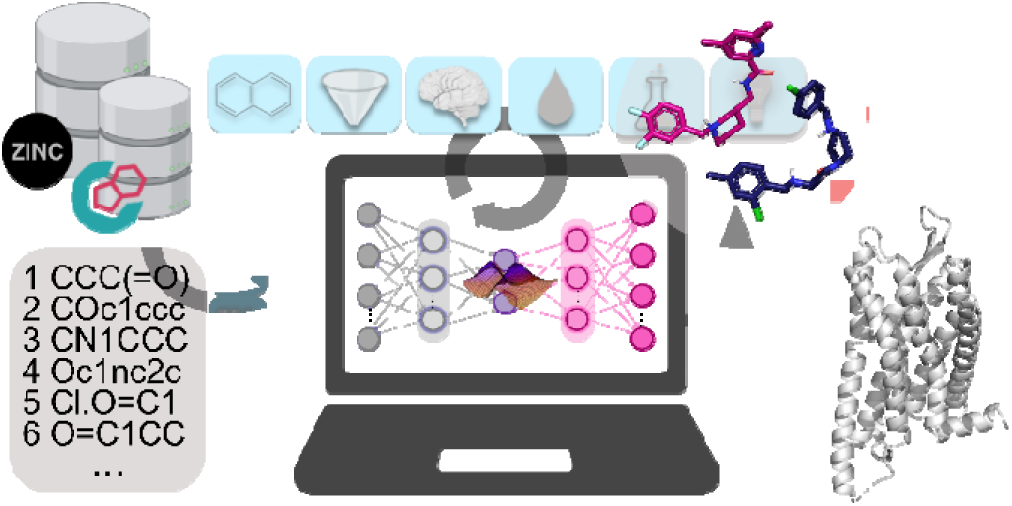

