## Supporting Information for "De Novo Design of κ-Opioid Receptor Antagonists Using a Generative Deep Learning Framework"

*[1]Department of Pharmacological Sciences, Icahn School of Medicine at Mount Sinai, New York, NY, USA; [2]National Institute of Mental Health, Psychoactive Drug Screening Program, Department of Pharmacology, University of North Carolina School of Medicine, Chapel Hill, NC, USA; [3]Mount Sinai Center for Therapeutics Discovery, Departments of Oncological Sciences and Neuroscience, Tisch Cancer Institute, Icahn School of Medicine at Mount Sinai, New York, NY, USA; [4]Division of Chemical Biology and Medicinal Chemistry, University of North Carolina at Chapel Hill Eshelman School of Pharmacy, Chapel Hill, NC, USA*

| | |
|---|---|
| Chemical synthesis of selected prioritized compounds | Pg. 1 |
| Table S1 | Pg. 8 |
| Table S2 | Pg. 9 |
| Table S3 | Pg. 10 |
| Figure S1 | Pg. 25 |
| Figure S2 | Pg. 26 |
| Figure S3 | Pg. 27 |
| Figure S4 | Pg. 28 |

**Chemical synthesis of selected prioritized compounds**

*Abbreviations:* 1-ethyl-3-(3-dimethylaminopropyl)carbodiimide (ECDI); 1-hydroxy-7-azabenzotriazole (HOAt); acetic acid (AcOH); dichloromethane (DCM); diisopropylethylamine (DIEA); dimethylformamide (DMF); ethyl acetate (EA); formaldehyde (HCHO); hexafluorophosphate azabenzotriazole tetramethyl uronium (HATU); high-performance liquid chromatography (HPLC); liquid chromatography–mass spectrometry (LCMS); methanol (MeOH); N-Methylmorpholine (NMM*);* sodium cyanoborohydride ($NaBH_3CN$); sodium iodide (NaI)*; s*odium sulfate ($Na_2SO_4$); trifluoracetic acid (TFA).

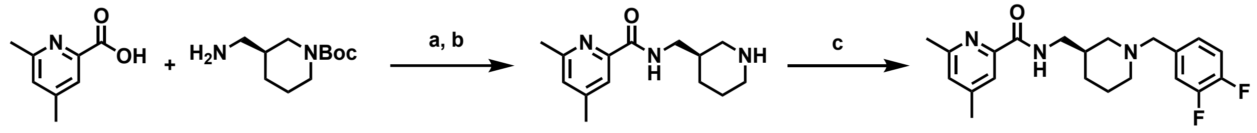


1 2 3 4

[a]Reagent and conditions: (a) HOAt, EDCI, NMM, DMF, rt; (b) TFA, DCM; (c) 4-(bromomethyl)-1,2-difluorobenzene, DIEA, DMF, rt;

*Scheme 1. Synthesis of compound FJRL-14 (4)*

*Synthesis of (R)-4,6-dimethyl-N-(piperidin-3-ylmethyl)picolinamide (3)*

To a solution of 4,6-dimethylpicolinic acid **1** (75.6 mg, 0.5 mmol) and tert-butyl (S)-3-(aminomethyl)piperidine-1-carboxylate **2** (107.1 mg, 0.5 mmol) in DMF (2 mL) was added HOAt (136.1 mg, 1 mmol), EDCI (191.7 mg, 1 mmol), and NMM (202.3mg, 2 mmol), and stirred at room temperature overnight. The reaction mixture was extracted with EA and the combined organic layer was washed with water, brine, dried by $Na_2SO_4$ and concentrated under vacuum. The residue was dissolved in DCM (3 mL), and TFA (1 mL) was added. After stirring at room temperature for 1 hour, the mixture was concentrated under vacuum and purified by column chromatography (C18, 10%-100% acetonitrile / 0.1% TFA in $H_2O$) to give compound **3** (78 mg, 63% yield in 2 steps) as yellow oil. LCMS calcd for $C_{14}H_{22}N_3O^+$ $[M + H]^+$ 248, found 248.

*Synthesis of (S)-N-((1-(3,4-difluorobenzyl)piperidin-3-yl)methyl)-4,6-dimethylpicolinamide (4)*

To a solution of (R)-4,6-dimethyl-N-(piperidin-3-ylmethyl)picolinamide **3** (49.5 mg, 0.2 mmol) in DMF (1 mL) was added 4-(bromomethyl)-1,2-difluorobenzene (41.4mg, 0.2 mmol) and DIEA (51.7 mg, 0.4 mmol), After being stirred at room temperature overnight, the resulting mixture was purified by preparative HPLC (10%-100% acetonitrile / 0.1% TFA in $H_2O$) to afford compound **4** as colorless oil (43 mg, 58% yield). $^1$H NMR (400 MHz, Methanol-$d_4$) δ 7.75 (s, 1H), 7.48 (t, $J$ = 9.8 Hz, 1H), 7.41 – 7.29 (m, 3H), 4.32 (s, 2H), 3.41 – 3.38 (m, 1H), 2.93 (dd, $J$ = 14.2, 11.1 Hz, 1H), 2.77 (t, $J$ = 12.3 Hz, 1H), 2.55 (s, 3H), 2.43 (s, 3H), 2.14 (s, 1H), 2.04 (d, $J$ = 14.5 Hz, 1H), 1.96 (d, $J$ = 13.5 Hz, 1H), 1.83 – 1.71 (m, 1H), 1.42 – 1.29 (m, 3H), 1.0 - 0.9 (m, 1H). LCMS calcd for $C_{21}H_{26}F_2N_3O^+$ $[M + H]^+$ 374, found 374.

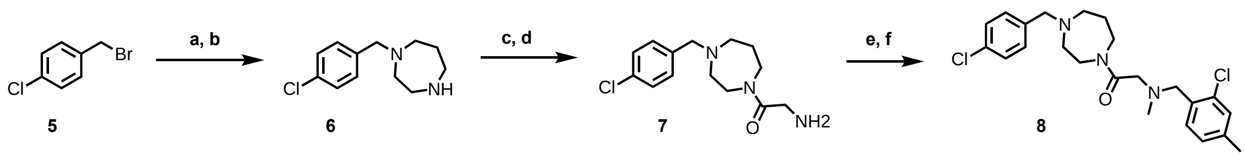

[a]Reagent and conditions: (a) *tert*-butyl 1,4-diazepane-1-carboxylate, DIEA, DMF, rt; (b) TFA, DCM; (c) (*tert*-butoxycarbonyl)glycine, HOAt, EDCI, NMM, DMF, rt; (d) TFA, DCM; (e) 2-chloro-4-methylbenzaldehyde, $NaBH_3CN$, MeOH, rt; (f) HCHO, rt;

*Scheme 2. Synthesis of compound FJRL-20 (8)*

*Synthesis of 1-(4-chlorobenzyl)-1,4-diazepane (6)*

To a solution of 1-(bromomethyl)-4-chlorobenzene **5** (114.6 mg, 1 mmol) in DMF (2 mL) was added *tert*-butyl 1,4-diazepane-1-carboxylate (200.3 mg, 1 mmol), DIEA (36.2 mg, 2 mmol) and stirred at room temperature overnight. The reaction mixture was extracted with EA and the combined organic layer was washed with water, brine, dried by $Na_2SO_4$ and concentrated under vacuum. The residue was dissolved in DCM (3 mL) and TFA (1 mL) was added. After stirring at room temperature for 1 hour, the mixture was concentrated under vacuum and purified by column chromatography (C18, 10%-100% acetonitrile / 0.1% TFA in $H_2O$) to give compound **6** (180 mg, 80% yield in 2 steps) as colorless oil. LCMS calcd for $C_{12}H_{18}ClN_2^+$ $[M + H]^+$ 225, found 225.

*Synthesis of 2-amino-1-(4-(4-chlorobenzyl)-1,4-diazepan-1-yl)ethan-1-one (7)*

To a solution of 1-(4-chlorobenzyl)-1,4-diazepane **6** (180 mg, 0.8 mmol) in DMF (2 mL) was added (*tert*-butoxycarbonyl)glycine (140.1mg, 0.8 mmol), was added HOAt (163.3 mg, 1.2 mmol), EDCI (230.0 mg, 1.2 mmol) and NMM (161.8 mg, 1.6 mmol) and stirred at room temperature overnight. The reaction mixture was extracted was extracted with EA and the combined organic layer was washed with water, brine, dried by $Na_2SO_4$ and concentrated under vacuum. The residue was dissolved in DCM (3 mL) and TFA (1 mL) was added. After stirring at room temperature for 1 hour, the mixture was concentrated under vacuum and purified by column chromatography (C18, 10%-100% acetonitrile / 0.1% TFA in $H_2O$) to give compound **7** (126 mg, 56% yield in 2 steps) as yellow oil. LCMS calcd for $C_{14}H_{21}ClN_3O^+$ $[M + H]^+$ 282, found 282.

*Synthesis of 2-((2-chloro-4-methylbenzyl)(methyl)amino)-1-(4-(4-chlorobenzyl)-1,4-diazepan-1-yl)ethan-1-one (8)*

To a solution of 2-amino-1-(4-(4-chlorobenzyl)-1,4-diazepan-1-yl)ethan-1-one **7** (42.3 mg, 0.15 mmol) in MeOH (2 mL) was added 2-chloro-4-methylbenzaldehyde (46.4 mg, 0.3 mmol), $NaBH_3CN$ (18.9 mg, 0.3 mmol) and AcOH (1 mL). After stirring at room temperature for 2 hour, HCHO (4.5 mg, 0.15 mmol) was added, After being stirred at room temperature overnight, the resulting mixture was purified by preparative HPLC (10%-100% acetonitrile / 0.1% TFA in $H_2O$) to afford compound **8** as colorless oil (22 mg, 34% yield in 2 steps). $^1H$ NMR (400 MHz, Methanold4) δ 7.65 – 7.52 (m, 5H), 7.43 (d, *J* = 12.3 Hz, 1H), 7.30 (dd, *J* = 17.3, 7.9 Hz, 1H), 4.49 – 4.38 (m, 4H), 4.36 (s, 1H), 3.62 (d, *J* = 16.0 Hz, 4H), 3.49 – 3.38 (m, 2H), 2.93 (d, *J* = 4.6 Hz, 2H), 2.61 (d, *J* = 5.3 Hz, 1H), 2.42 (s, 3H), 2.39 (s, 3H), 2.34 (s, 1H), 2.27 (s, 1H). LCMS calcd for $C_{23}H_{30}Cl_2N_3O^+$ $[M + H]^+$ 434, found 434.

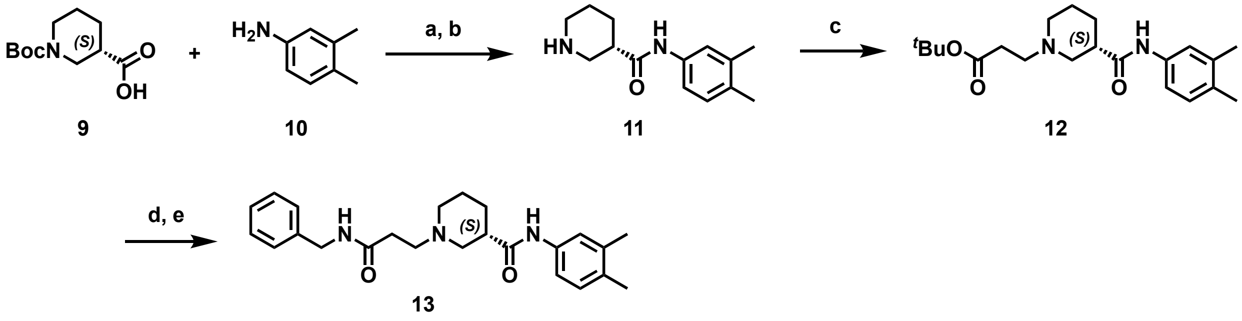


[a]Reagent and conditions: (a) HOAt, EDCI, NMM, DMF, rt; (b) TFA, DCM; (c) *tert*-butyl 3-bromopropanoate, DIEA, NaI, DMF, 50°C; (d) TFA, DCM; (e) Benzylamine, HOAt, EDCI, NMM, DMF, rt;

*Scheme 3. Synthesis of compound FJRL-29 (13)*

*Synthesis of (S)-N-(3,4-dimethylphenyl)piperidine-3-carboxamide (11)*

To a solution of (S)-1-(*tert*-butoxycarbonyl)piperidine-3-carboxylic acid **9** (114.6 mg, 0.5 mmol) and 3,4-dimethylaniline **10** (60.6 mg, 0.5 mmol) in DMF (2 mL) was added HOAt (136.1 mg, 1 mmol) EDCI (191.7 mg, 1 mmol) and NMM (202.3mg, 2 mmol) and stirred at room temperature overnight. The reaction mixture was extracted with EA and the combined organic layer was washed with water, brine, dried by $Na_2SO_4$ and concentrated under vacuum. The residue was dissolved in DCM (3 mL) and TFA (1 mL) was added. After stirring at room temperature for 1 hour, the mixture was concentrated under vacuum and purified by column chromatography (C18, 10%-100% acetonitrile / 0.1% TFA in $H_2O$) to give compound **11** (51 mg, 44% yield in 2 steps) as yellow oil. LCMS calcd for $C_{14}H_{21}N_2O^+$ $[M + H]^+$ 233, found 233.

*Synthesis of tert-butyl (S)-3-(3-((3,4-dimethylphenyl)carbamoyl)piperidin-1-yl)propanoate (12)*

To a solution of (S)-N-(3,4-dimethylphenyl)piperidine-3-carboxamide **11** (46.5 mg, 0.2 mmol) in DMF (1 mL) was added *tert*-butyl 3-bromopropanoate (125.4 mg, 0.6 mmol), DIEA (77.5 mg, 0.6 mmol) and NaI (89.9 mg, 0.6 mmol) and stirred at 50 °C for 6 h. The mixture was purified by column chromatography (C18, 10%-100% acetonitrile / 0.1% TFA in $H_2O$) to give compound **12** (48 mg, 66% yield in 2 steps) as yellow oil. LCMS calcd for $C_{21}H_{33}N_2O_3^+$ $[M + H]^+$ 361, found 361.

*Synthesis of (S)-N-((1-(3,4-difluorobenzyl)piperidin-3-yl)methyl)-4,6-dimethylpicolinamide (13)*

To a solution of *tert*-butyl (S)-3-(3-((3,4-dimethylphenyl)carbamoyl)piperidin-1-yl)propanoate **12** (36.0 mg, 0.1 mmol) in DCM (2 mL) was added TFA (1 mL). After stirring at room temperature for 1 hour, the mixture was concentrated under vacuum. The residue was dissolved in DMF (1 mL), then Benzylamine (10.7 mg, 0.1 mmol), HOAt (27.2 mg, 0.2 mmol), EDCI (38.3 mg, 0.2 mmol) and NMM (40.5 mg, 0.4 mmol) were added. After being stirred at room temperature overnight, the resulting mixture was purified by preparative HPLC (10%-100% acetonitrile / 0.1% TFA in $H_2O$) to afford compound **13** as colorless oil (24 mg, 61% yield). $^1H$ NMR (400 MHz, Methanol-$d_4$) δ 7.33 – 7.13 (m, 7H), 7.01 (dd, *J* = 8.7, 3.2 Hz, 1H), 4.39 – 4.31 (m, 2H), 3.71 (d, *J* = 12.1 Hz, 1H), 3.56 (d, *J* = 12.2 Hz, 1H), 3.41 (d, *J* = 7.2 Hz, 1H), 3.34 (q, *J* = 6.3 Hz, 1H), 3.50 – 2.90 (m, 2H), 2.83 – 2.69 (m, 3H), 2.17 (d, *J* = 5.8 Hz, 6H), 2.04 (d, *J* = 14.0 Hz, 2H), 1.98 – 1.85 (m, 2H). LCMS calcd for $C_{24}H_{32}N_3O_2^+$ $[M + H]^+$ 394, found 394.

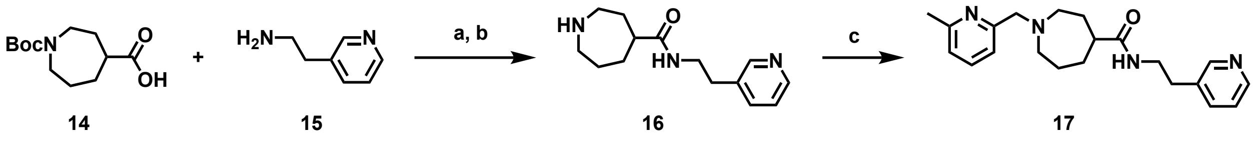

[a]Reagent and conditions: (a) HOAt, EDCI, NMM, DMF, rt; (b) TFA, DCM; (c) 2-(bromomethyl)-6-methylpyridine, DIEA, DMF, rt;

*Scheme 4. Synthesis of compound FJRL-36 (17)*

*Synthesis of N-(2-(pyridin-3-yl)ethyl)azepane-4-carboxamide (16)*

To a solution of 1-(tert-butoxycarbonyl)azepane-4-carboxylic acid **14** (48.7 mg, 0.2 mmol) and 2-(pyridin-3-yl)ethan-1-amine **15** (31.7 mg, 0.2 mmol) in DMF (3 mL) was added HOAt (54.4 mg, 0.4 mmol), EDCI (76.7 mg, 0.4 mmol) and NMM (80.9 mg, 0.8 mmol) and stirred at room temperature overnight. The reaction mixture was extracted with EA and the combined organic layer was washed with water, brine, dried by $Na_2SO_4$ and concentrated under vacuum. The residue was dissolved in DCM (3 mL) and TFA (1 mL) was added. After stirring at room temperature for 1 hour, the mixture was concentrated under vacuum and purified by column chromatography (C18, 10%-100% acetonitrile / 0.1% TFA in $H_2O$) to give compound **16** (35 mg, 71% yield in 2 steps) as yellow oil. LCMS calcd for $C_{14}H_{22}N_3O^+$ $[M + H]^+$ 248, found 248.

*Synthesis of 1-((6-methylpyridin-2-yl)methyl)-N-(2-(pyridin-3-yl)ethyl)azepane-4-carboxamide (17)*

To a solution of N-(2-(pyridin-3-yl)ethyl)azepane-4-carboxamide **16** (34.6 mg, 0.14 mmol) in DMF (1 mL) was added 2-(bromomethyl)-6-methylpyridine (26.0 mg, 0.14 mmol) and DIEA (36.2 mg, 0.28 mmol), After being stirred at room temperature overnight, the resulting mixture was purified by preparative HPLC (10%-100% acetonitrile / 0.1% TFA in $H_2O$) to afford compound **17** as colorless oil (36 mg, 73% yield). $^1H$ NMR (400 MHz, Methanol-$d_4$) δ 8.72 (d, *J*

= 18.6 Hz, 2H), 8.41 (d, *J* = 8.1 Hz, 1H), 7.93 (dd, *J* = 8.0, 5.6 Hz, 1H), 7.79 (t, *J* = 7.7 Hz, 1H), 7.31 (dd, *J* = 17.3, 7.7 Hz, 2H), 4.48 (s, 2H), 3.61 – 3.51 (m, 2H), 3.05 (t, *J* = 6.9 Hz, 2H), 2.60 (s, 3H), 2.14 – 1.93 (m, 4H), 1.87 – 1.77 (m, 1H), 1.39 (dd, *J* = 6.8, 3.6 Hz, 6H). LCMS calcd for $C_{21}H_{29}N_4O^+$ $[M + H]^+$ 353, found 353.

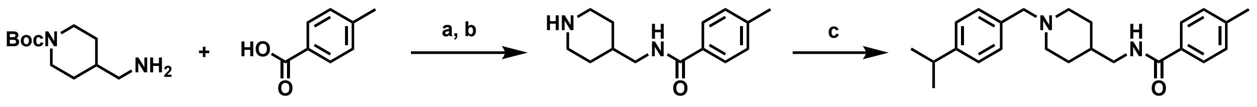


18 19 20 21

[a]Reagent and conditions: (a) HATU, DIEA, DMF, rt; (b) TFA, DCM; (c) 4-isopropylbenzaldehyde, $NaBH_3CN$, MeOH, rt

*Scheme 5. Synthesis of compound FJRL-91 (21)*

*Synthesis of 4-methyl-N-(piperidin-4-ylmethyl)benzamide (20)*

To a solution of *tert*-butyl 4-(aminomethyl)piperidine-1-carboxylate **18** (214.3 mg, 1 mmol) and 4-methylbenzoic acid **19** (136.2 mg, 1 mmol) in DMF (3 mL) was added HATU (456.3, 1.2 mmol) and DIEA (258.5 mg, 2 mmol) and stirred at room temperature overnight. The reaction mixture was extracted with EA and the combined organic layer was washed with water, brine, dried by $Na_2SO_4$ and concentrated under vacuum. The residue was dissolved in DCM (3 mL) and TFA (1 mL) was added. After stirring at room temperature for 1 hour, the mixture was concentrated under vacuum and purified by column chromatography (C18, 10%-100% acetonitrile / 0.1% TFA in $H_2O$) to give compound **20** (135 mg, 58% yield in 2 steps) as yellow oil. LCMS calcd for $C_{14}H_{21}N_2O^+$ $[M + H]^+$ 233, found 233.

*Synthesis of N-((1-(4-isopropylbenzyl)piperidin-4-yl)methyl)-4-methylbenzamide (21)*

To a solution of 4-methyl-N-(piperidin-4-ylmethyl)benzamide **20** (44.5 mg, 0.3 mmol) in MeOH (1 mL) was added 4-isopropylbenzaldehyde (69.7 mg, 0.3 mmol) and $NaBH_3CN$ (37.8 mg, 0.6 mmol). After being stirred at room temperature overnight, the resulting mixture was purified by preparative HPLC (10%-100% acetonitrile / 0.1% TFA in $H_2O$) to afford compound **21** as white solid (56 mg, 51% yield). $^1$H NMR (400 MHz, Methanol-$d_4$) δ 7.60 (d, *J* = 8.2 Hz, 2H), 7.34 – 7.23 (m, 4H), 7.17 (d, *J* = 7.9 Hz, 2H), 4.15 (s, 2H), 3.45 – 3.36 (m, 2H), 3.19 (d, *J* = 5.8 Hz, 2H), 2.93 – 2.81 (m, 3H), 2.29 (s, 3H), 1.96 – 1.78 (mz, 3H), 1.48 – 1.33 (m, 2H), 1.16 (d, *J* = 6.9 Hz, 6H). LCMS calcd for $C_{24}H_{33}N_2O^+$ $[M + H]^+$ 365, found 365.

**Table S1. Performance of the deep generative model using reference datasets after pre-training on Dataset #1.** FCD: Fréchet ChemNet Distance; Novelty: fraction of generated molecules not present in the training set; Frag: similarity of distribution of BRICS fragments; Scaff: similarity of distribution of Bemis-Murko scaffolds; SNN: average Tanimoto similarity of Morgan fingerprints to the nearest neighbor. An epoch consists of one pass over the entire Dataset #1.

| # Epochs | FCD Test | FCD Test Scaffolds | Novelty | Frag Test | Frag Test Scaffolds | Scaff Test | Scaff Test Scaffolds | SNN Test | SNN Test Scaffolds |
|---|---|---|---|---|---|---|---|---|---|
| 0 | 49.66 | 49.70 | 1.0 | 1.77 | 1.72 | NaN | NaN | 0.09 | 0.08 |
| 2 | 3.71 | 4.21 | 0.99 | 0.98 | 0.98 | 0.67 | 0.12 | 0.45 | 0.43 |
| 4 | 3.88 | 4.30 | 1.0 | 0.98 | 0.98 | 0.57 | 0.13 | 0.44 | 0.42 |
| 6 | 3.84 | 4.20 | 0.99 | 0.98 | 0.98 | 0.58 | 0.12 | 0.45 | 0.43 |
| 8 | 4.39 | 4.82 | 0.99 | 0.98 | 0.97 | 0.64 | 0.13 | 0.44 | 0.42 |
| 10 | 3.86 | 4.31 | 0.99 | 0.98 | 0.98 | 0.63 | 0.11 | 0.44 | 0.42 |
| 15 | 3.69 | 3.98 | 1.0 | 0.99 | 0.98 | 0.60 | 0.10 | 0.44 | 0.42 |
| 20 | 4.18 | 4.79 | 0.99 | 0.98 | 0.98 | 0.68 | 0.12 | 0.45 | 0.43 |
| 25 | 4.07 | 4.40 | 0.99 | 0.99 | 0.98 | 0.59 | 0.10 | 0.44 | 0.42 |

**Table S2. Performance of the deep generative model using reference datasets after pre-training with Dataset #2.** FCD: Fréchet ChemNet Distance; Novelty: fraction of generated molecules not present in the training set; Frag: similarity of distribution of BRICS fragments; Scaff: similarity of distribution of Bermis-Murko scaffolds; SNN: average Tanimoto similarity of Morgan fingerprints to the nearest neighbor. An update consists of a pass over one batch from Dataset #2, composed of 100 molecules from Dataset #1, 50 molecules from the positive KOR inhibitor ($IC_{50}$ < 1 μM) dataset, and 50 molecules from the negative KOR ($IC_{50}$ ≥ 1μM) inhibitor dataset.

| # Updates | FCD Test | FCD Test Scaffolds | Novelty | Frag Test | Frag Test Scaffolds | Scaff Test | Scaff Test Scaffolds | SNN Test | SNN Test Scaffolds |
|---|---|---|---|---|---|---|---|---|---|
| 0 | 4.18 | 4.79 | 0.99 | 0.98 | 0.98 | 0.68 | 0.12 | 0.45 | 0.43 |
| 300,000 | 4.02 | 4.56 | 0.99 | 0.98 | 0.98 | 0.65 | 0.11 | 0.47 | 0.40 |

**Table S3. The 169 molecules prioritized for chemical synthesis on the basis of their similarity to JDTic's interactions with KOR.** The compounds are sorted by decreasing similarity to JDTic's interaction fingerprints (SIFt Tc). The method used to assess pharmacophore similarity (Pharm) during reinforcement learning is reported for each molecule and the five compounds synthesized for experimental testing are indicated with red boxes.

| | | |
|---|---|---|
| 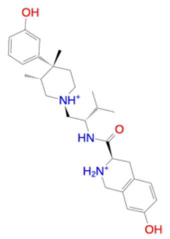 | 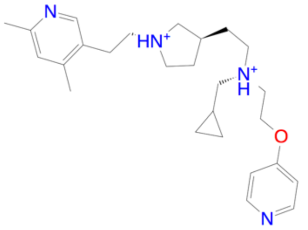 | 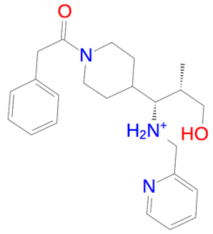 |
| Name JDTic<br>SIFt Tc 1.0<br>Pharm assessed by None | Name FJRL-1<br>SIFt Tc 0.905<br>Pharm assessed by IChem | Name FJRL-2<br>SIFt Tc 0.895<br>Pharm assessed by USRCAT |
| 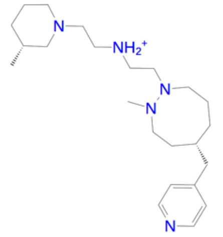 | 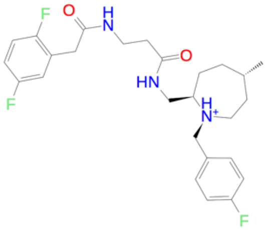 | 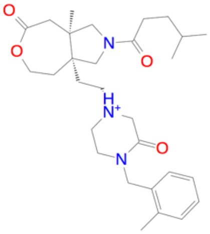 |
| Name FJRL-3<br>SIFt Tc 0.895<br>Pharm assessed by IChem | Name FJRL-4<br>SIFt Tc 0.857<br>Pharm assessed by USRCAT | Name FJRL-5<br>SIFt Tc 0.857<br>Pharm assessed by IChem |
| 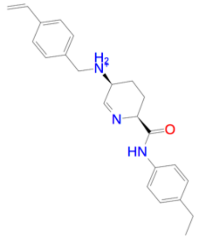 | 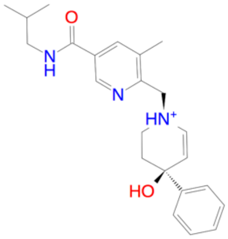 | 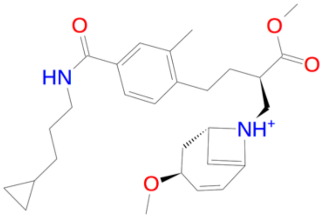 |
| Name FJRL-6<br>SIFt Tc 0.857<br>Pharm assessed by IChem | Name FJRL-7<br>SIFt Tc 0.857<br>Pharm assessed by IChem | Name FJRL-8<br>SIFt Tc 0.857<br>Pharm assessed by IChem |
| 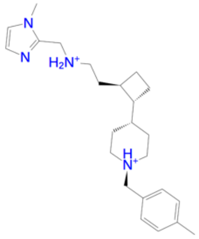 | 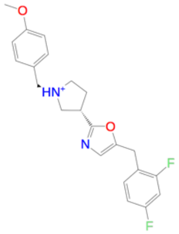 | 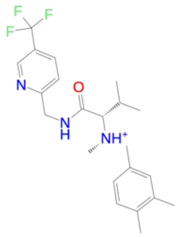 |
| Name FJRL-9<br>SIFt Tc 0.85<br>Pharm assessed by IChem | Name FJRL-10<br>SIFt Tc 0.85<br>Pharm assessed by IChem | Name FJRL-11<br>SIFt Tc 0.85<br>Pharm assessed by IChem |

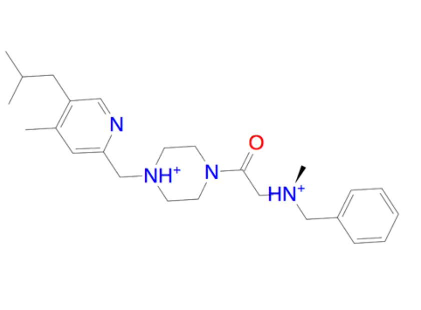

| | |
|---|---|
| Name | FJRL-12 |
| SIFt Tc | 0.85 |
| Pharm assessed by | IChem |

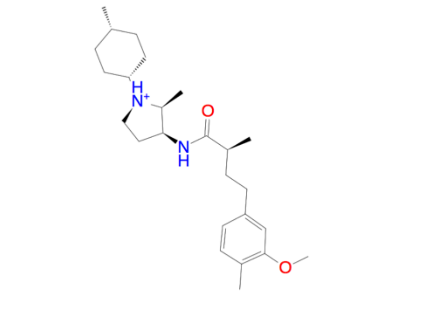

| | |
|---|---|
| Name | FJRL-13 |
| SIFt Tc | 0.85 |
| Pharm assessed by | IChem |

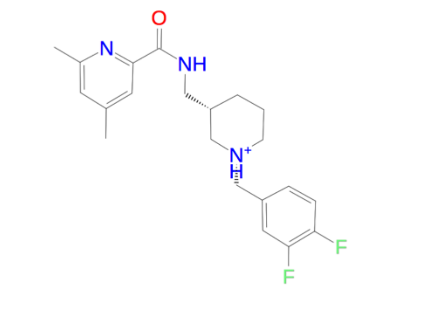

| | |
|---|---|
| Name | FJRL-14 |
| SIFt Tc | 0.842 |
| Pharm assessed by | USRCAT |

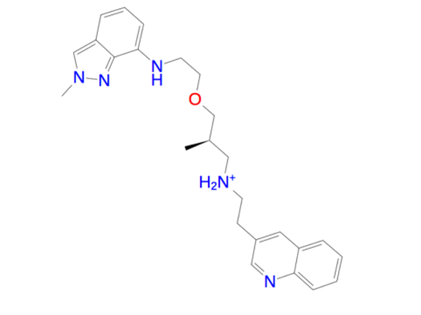

| | |
|---|---|
| Name | FJRL-15 |
| SIFt Tc | 0.826 |
| Pharm assessed by | USRCAT |

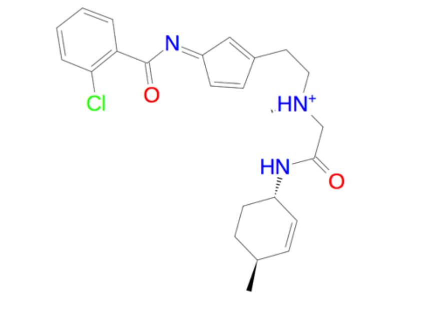

| | |
|---|---|
| Name | FJRL-16 |
| SIFt Tc | 0.818 |
| Pharm assessed by | IChem |

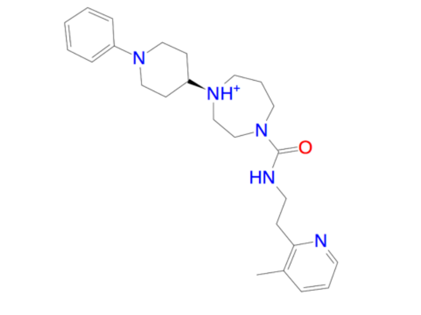

| | |
|---|---|
| Name | FJRL-17 |
| SIFt Tc | 0.818 |
| Pharm assessed by | IChem |

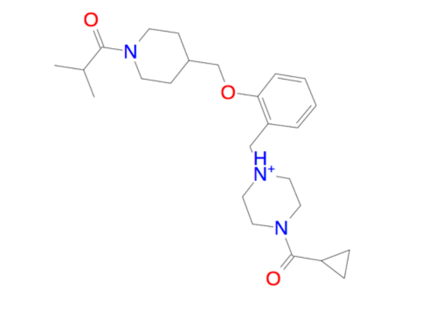

| | |
|---|---|
| Name | FJRL-18 |
| SIFt Tc | 0.81 |
| Pharm assessed by | IChem |

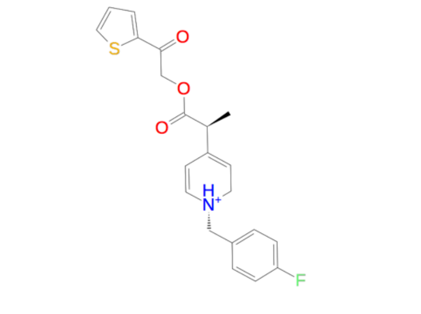

| | |
|---|---|
| Name | FJRL-19 |
| SIFt Tc | 0.81 |
| Pharm assessed by | IChem |

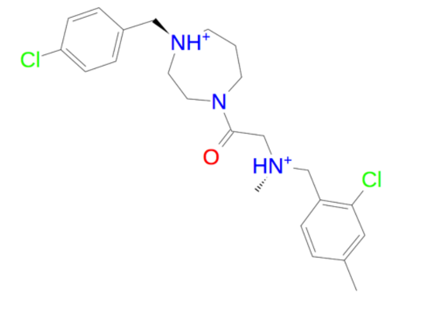

| | |
|---|---|
| Name | FJRL-20 |
| SIFt Tc | 0.81 |
| Pharm assessed by | IChem |

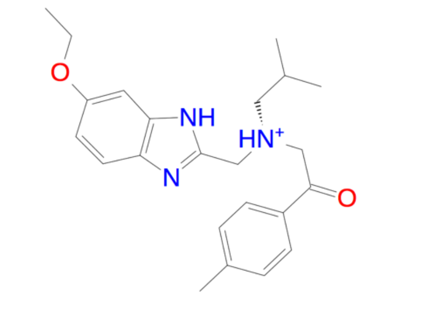

| | |
|---|---|
| Name | FJRL-21 |
| SIFt Tc | 0.81 |
| Pharm assessed by | IChem |

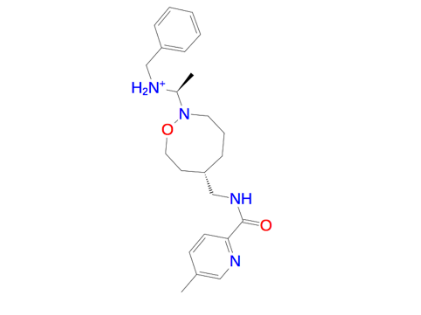

| | |
|---|---|
| Name | FJRL-22 |
| SIFt Tc | 0.81 |
| Pharm assessed by | USRCAT |

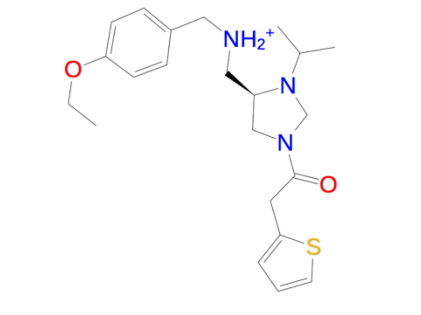

| | |
|---|---|
| Name | FJRL-23 |
| SIFt Tc | 0.81 |
| Pharm assessed by | IChem |

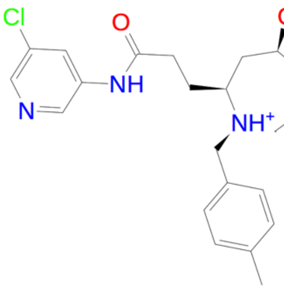

| | |
|---|---|
| Name | FJRL-24 |
| SIFt Tc | 0.81 |
| Pharm assessed by | IChem |

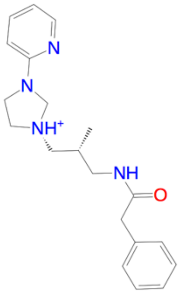

| | |
|---|---|
| Name | FJRL-25 |
| SIFt Tc | 0.81 |
| Pharm assessed by | USRCAT |

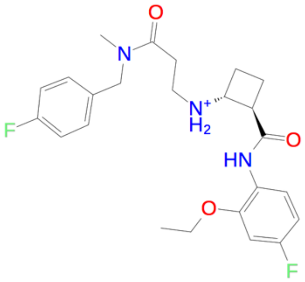

| | |
|---|---|
| Name | FJRL-26 |
| SIFt Tc | 0.81 |
| Pharm assessed by | USRCAT |

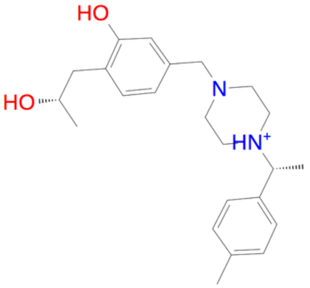

| | |
|---|---|
| Name | FJRL-27 |
| SIFt Tc | 0.81 |
| Pharm assessed by | USRCAT |

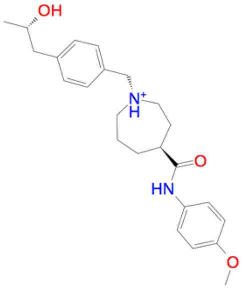

| | |
|---|---|
| Name | FJRL-28 |
| SIFt Tc | 0.81 |
| Pharm assessed by | USRCAT |

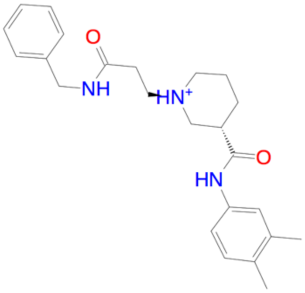

| | |
|---|---|
| Name | FJRL-29 |
| SIFt Tc | 0.81 |
| Pharm assessed by | IChem |

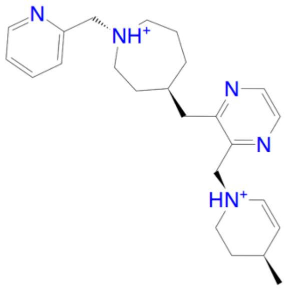

| | |
|---|---|
| Name | FJRL-30 |
| SIFt Tc | 0.81 |
| Pharm assessed by | USRCAT |

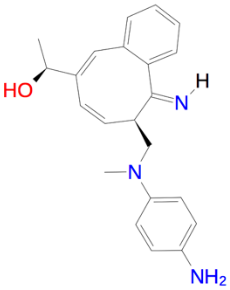

| | |
|---|---|
| Name | FJRL-31 |
| SIFt Tc | 0.81 |
| Pharm assessed by | USRCAT |

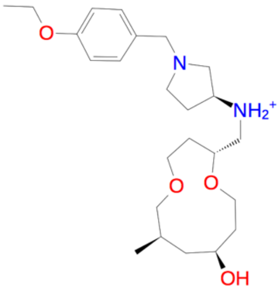

| | |
|---|---|
| Name | FJRL-32 |
| SIFt Tc | 0.81 |
| Pharm assessed by | IChem |

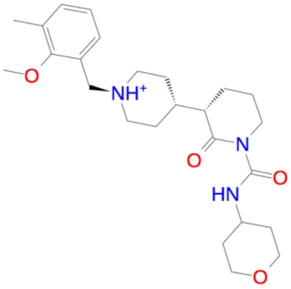

| | |
|---|---|
| Name | FJRL-33 |
| SIFt Tc | 0.81 |
| Pharm assessed by | IChem |

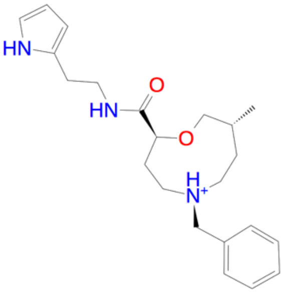

| | |
|---|---|
| Name | FJRL-34 |
| SIFt Tc | 0.81 |
| Pharm assessed by | USRCAT |

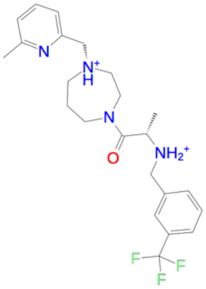

| | |
|---|---|
| Name | FJRL-35 |
| SIFt Tc | 0.81 |
| Pharm assessed by | IChem |

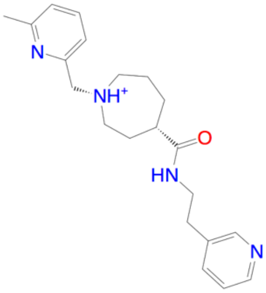

| | |
|---|---|
| Name | FJRL-36 |
| SIFt Tc | 0.81 |
| Pharm assessed by | USRCAT |

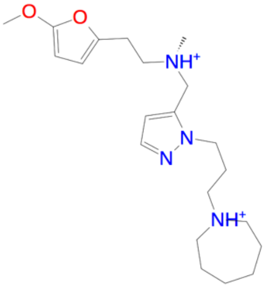

| | |
|---|---|
| Name | FJRL-37 |
| SIFt Tc | 0.81 |
| Pharm assessed by | USRCAT |

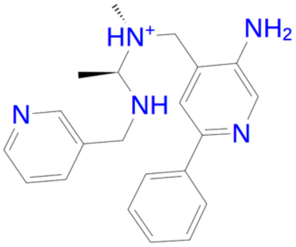

| | |
|---|---|
| Name | FJRL-38 |
| SIFt Tc | 0.81 |
| Pharm assessed by | USRCAT |

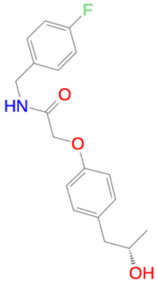

| | |
|---|---|
| Name | FJRL-39 |
| SIFt Tc | 0.8 |
| Pharm assessed by | USRCAT |

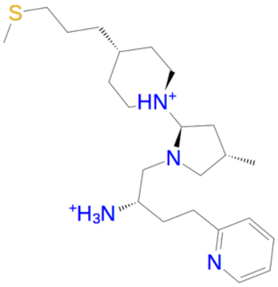

| | |
|---|---|
| Name | FJRL-40 |
| SIFt Tc | 0.8 |
| Pharm assessed by | IChem |

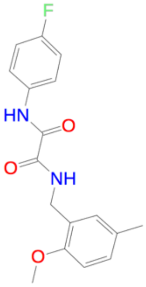

| | |
|---|---|
| Name | FJRL-41 |
| SIFt Tc | 0.8 |
| Pharm assessed by | USRCAT |

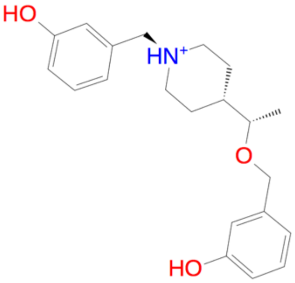

| | |
|---|---|
| Name | FJRL-42 |
| SIFt Tc | 0.8 |
| Pharm assessed by | USRCAT |

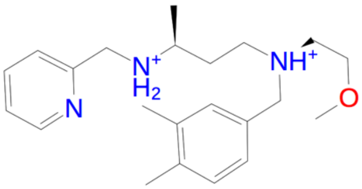

| | |
|---|---|
| Name | FJRL-43 |
| SIFt Tc | 0.8 |
| Pharm assessed by | USRCAT |

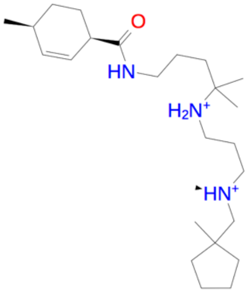

| | |
|---|---|
| Name | FJRL-44 |
| SIFt Tc | 0.8 |
| Pharm assessed by | IChem |

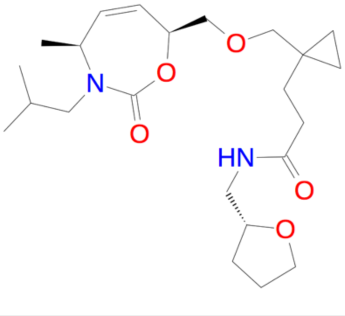

| | |
|---|---|
| Name | FJRL-45 |
| SIFt Tc | 0.8 |
| Pharm assessed by | IChem |

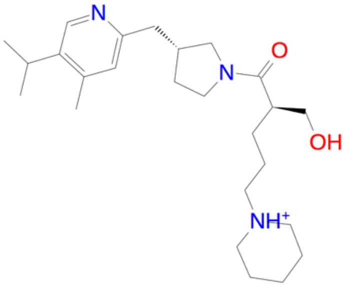

| | |
|---|---|
| Name | FJRL-46 |
| SIFt Tc | 0.8 |
| Pharm assessed by | IChem |

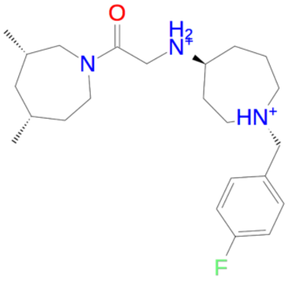

| | |
|---|---|
| Name | FJRL-47 |
| SIFt Tc | 0.8 |
| Pharm assessed by | IChem |

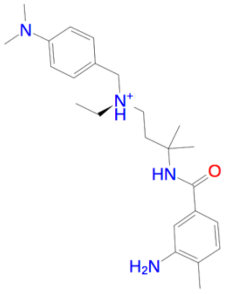

| | |
|---|---|
| Name | FJRL-48 |
| SIFt Tc | 0.8 |
| Pharm assessed by | IChem |

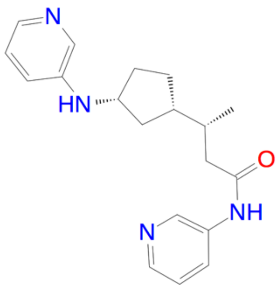

| | |
|---|---|
| Name | FJRL-49 |
| SIFt Tc | 0.8 |
| Pharm assessed by | USRCAT |

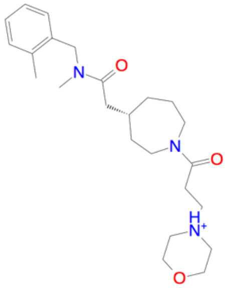

| | |
|---|---|
| Name | FJRL-50 |
| SIFt Tc | 0.8 |
| Pharm assessed by | IChem |

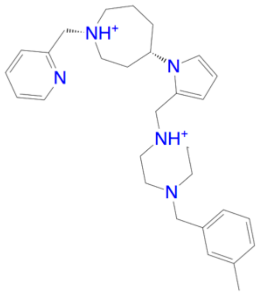

| | |
|---|---|
| Name | FJRL-51 |
| SIFt Tc | 0.792 |
| Pharm assessed by | IChem |

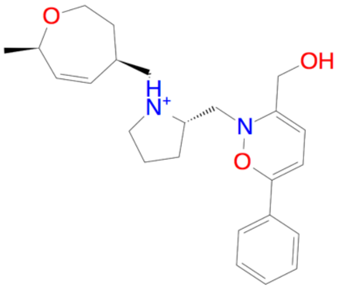

| | |
|---|---|
| Name | FJRL-52 |
| SIFt Tc | 0.789 |
| Pharm assessed by | IChem |

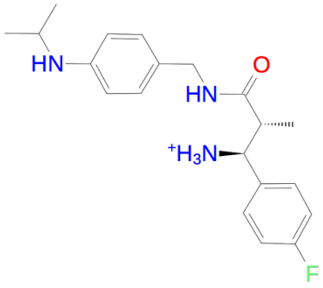

| | |
|---|---|
| Name | FJRL-53 |
| SIFt Tc | 0.789 |
| Pharm assessed by | USRCAT |

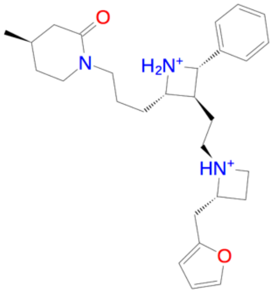

| | |
|---|---|
| Name | FJRL-54 |
| SIFt Tc | 0.783 |
| Pharm assessed by | IChem |

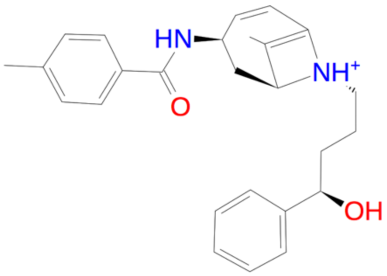

| | |
|---|---|
| Name | FJRL-55 |
| SIFt Tc | 0.783 |
| Pharm assessed by | USRCAT |

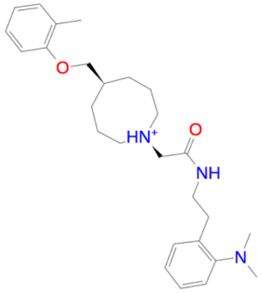

| | |
|---|---|
| Name | FJRL-56 |
| SIFt Tc | 0.783 |
| Pharm assessed by | IChem |

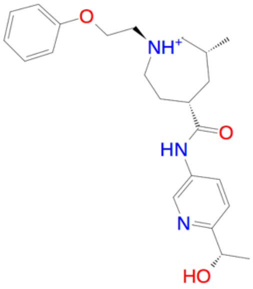

| | |
|---|---|
| Name | FJRL-57 |
| SIFt Tc | 0.783 |
| Pharm assessed by | USRCAT |

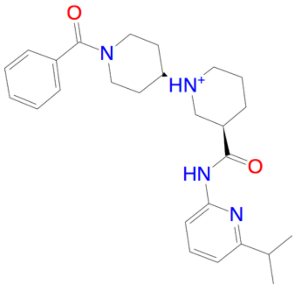

| | |
|---|---|
| Name | FJRL-58 |
| SIFt Tc | 0.783 |
| Pharm assessed by | IChem |

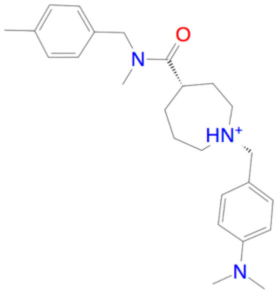

| | |
|---|---|
| Name | FJRL-59 |
| SIFt Tc | 0.783 |
| Pharm assessed by | IChem |

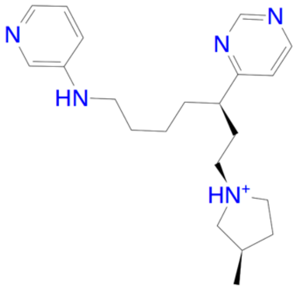

| | |
|---|---|
| Name | FJRL-60 |
| SIFt Tc | 0.783 |
| Pharm assessed by | USRCAT |

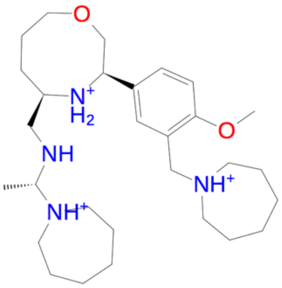

| | |
|---|---|
| Name | FJRL-61 |
| SIFt Tc | 0.783 |
| Pharm assessed by | IChem |

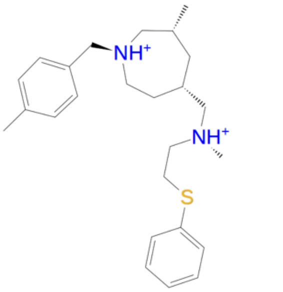

| | |
|---|---|
| Name | FJRL-62 |
| SIFt Tc | 0.783 |
| Pharm assessed by | IChem |

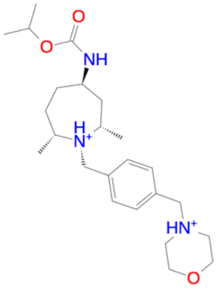

| | |
|---|---|
| Name | FJRL-63 |
| SIFt Tc | 0.783 |
| Pharm assessed by | IChem |

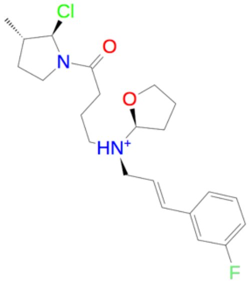

| | |
|---|---|
| Name | FJRL-64 |
| SIFt Tc | 0.773 |
| Pharm assessed by | IChem |

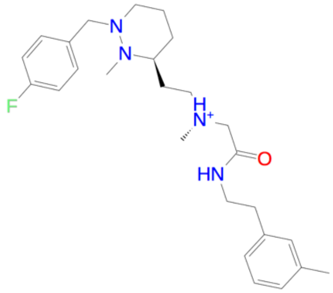

| | |
|---|---|
| Name | FJRL-65 |
| SIFt Tc | 0.773 |
| Pharm assessed by | IChem |

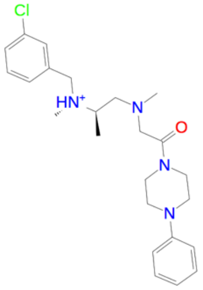

| | |
|---|---|
| Name | FJRL-66 |
| SIFt Tc | 0.773 |
| Pharm assessed by | IChem |

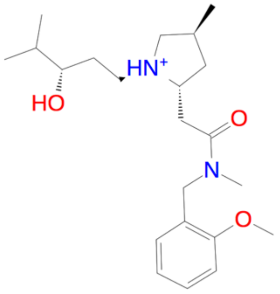

| | |
|---|---|
| Name | FJRL-67 |
| SIFt Tc | 0.773 |
| Pharm assessed by | IChem |

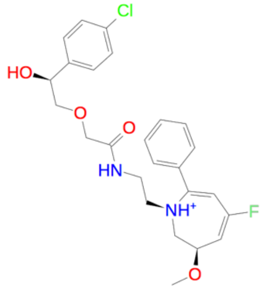

| | |
|---|---|
| Name | FJRL-68 |
| SIFt Tc | 0.773 |
| Pharm assessed by | IChem |

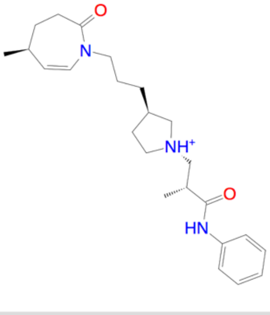

| | |
|---|---|
| Name | FJRL-69 |
| SIFt Tc | 0.773 |
| Pharm assessed by | IChem |

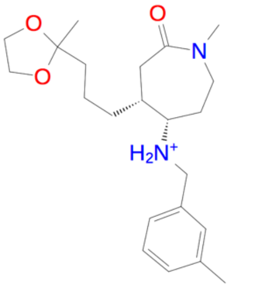

| | |
|---|---|
| Name | FJRL-70 |
| SIFt Tc | 0.773 |
| Pharm assessed by | IChem |

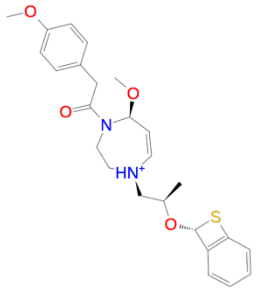

| | |
|---|---|
| Name | FJRL-71 |
| SIFt Tc | 0.773 |
| Pharm assessed by | IChem |

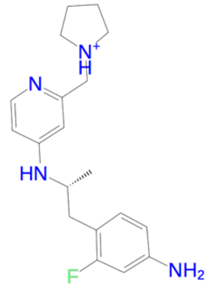

| | |
|---|---|
| Name | FJRL-72 |
| SIFt Tc | 0.773 |
| Pharm assessed by | USRCAT |

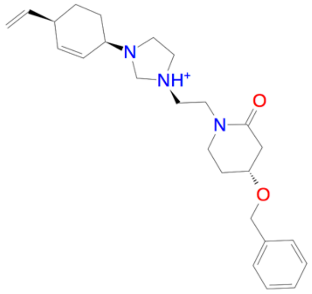

| | |
|---|---|
| Name | FJRL-73 |
| SIFt Tc | 0.773 |
| Pharm assessed by | IChem |

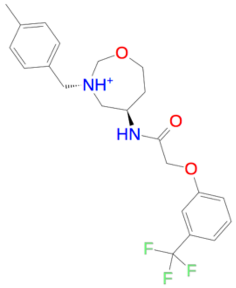

| | |
|---|---|
| Name | FJRL-74 |
| SIFt Tc | 0.773 |
| Pharm assessed by | IChem |

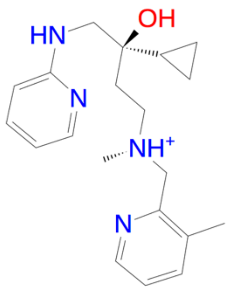

| | |
|---|---|
| Name | FJRL-75 |
| SIFt Tc | 0.773 |
| Pharm assessed by | USRCAT |

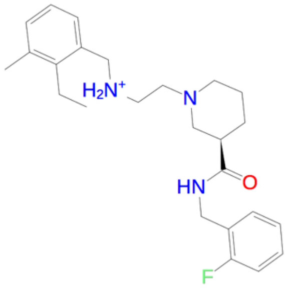

| | |
|---|---|
| Name | FJRL-76 |
| SIFt Tc | 0.773 |
| Pharm assessed by | IChem |

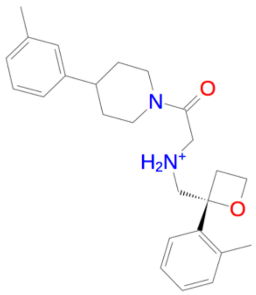

| | |
|---|---|
| Name | FJRL-77 |
| SIFt Tc | 0.773 |
| Pharm assessed by | IChem |

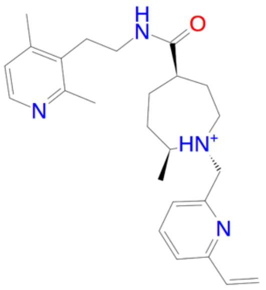

| | |
|---|---|
| Name | FJRL-78 |
| SIFt Tc | 0.773 |
| Pharm assessed by | USRCAT |

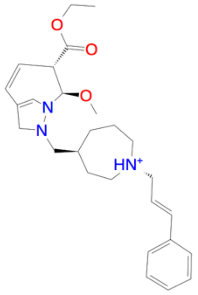

| | |
|---|---|
| Name | FJRL-79 |
| SIFt Tc | 0.773 |
| Pharm assessed by | IChem |

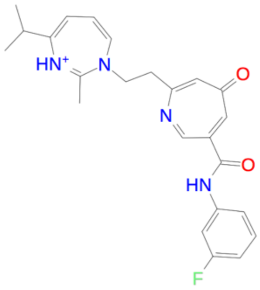

| | |
|---|---|
| Name | FJRL-80 |
| SIFt Tc | 0.773 |
| Pharm assessed by | USRCAT |

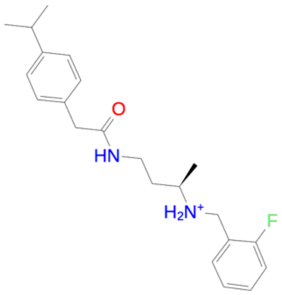

| | |
|---|---|
| Name | FJRL-81 |
| SIFt Tc | 0.773 |
| Pharm assessed by | USRCAT |

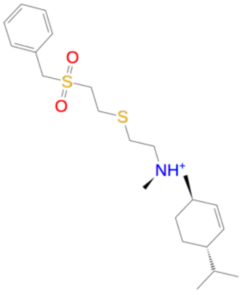

| | |
|---|---|
| Name | FJRL-82 |
| SIFt Tc | 0.773 |
| Pharm assessed by | IChem |

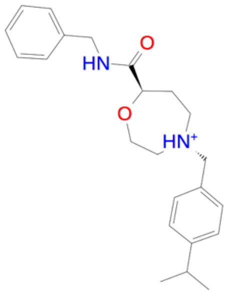

| | |
|---|---|
| Name | FJRL-83 |
| SIFt Tc | 0.773 |
| Pharm assessed by | IChem |

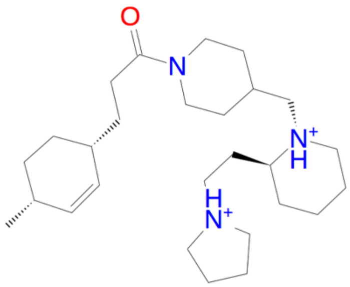

| | |
|---|---|
| Name | FJRL-84 |
| SIFt Tc | 0.773 |
| Pharm assessed by | IChem |

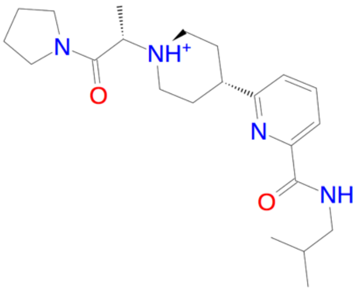

| | |
|---|---|
| Name | FJRL-85 |
| SIFt Tc | 0.773 |
| Pharm assessed by | IChem |

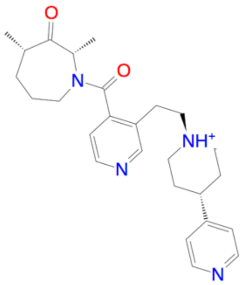

| | |
|---|---|
| Name | FJRL-86 |
| SIFt Tc | 0.773 |
| Pharm assessed by | IChem |

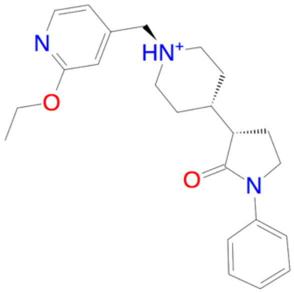

| | |
|---|---|
| Name | FJRL-87 |
| SIFt Tc | 0.773 |
| Pharm assessed by | IChem |

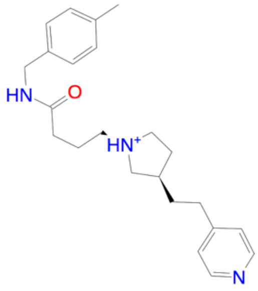

| | |
|---|---|
| Name | FJRL-88 |
| SIFt Tc | 0.773 |
| Pharm assessed by | IChem |

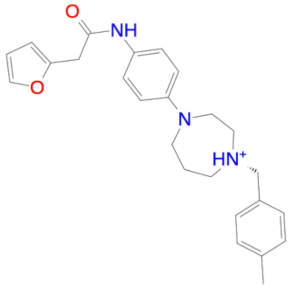

| | |
|---|---|
| Name | FJRL-89 |
| SIFt Tc | 0.773 |
| Pharm assessed by | IChem |

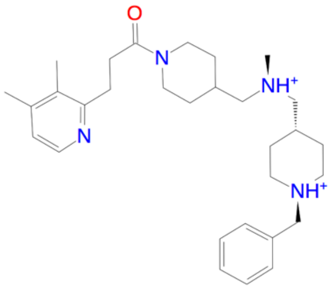

| | |
|---|---|
| Name | FJRL-90 |
| SIFt Tc | 0.773 |
| Pharm assessed by | IChem |

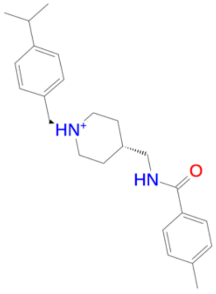

| | |
|---|---|
| Name | FJRL-91 |
| SIFt Tc | 0.773 |
| Pharm assessed by | IChem |

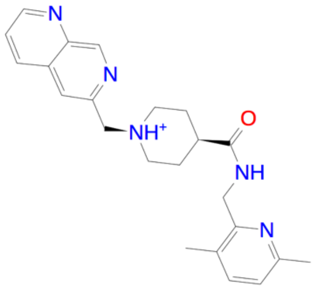

| | |
|---|---|
| Name | FJRL-92 |
| SIFt Tc | 0.773 |
| Pharm assessed by | USRCAT |

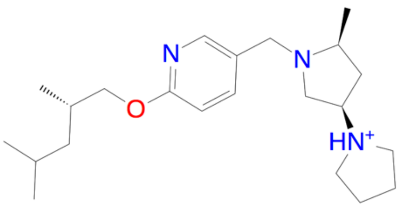

| | |
|---|---|
| Name | FJRL-93 |
| SIFt Tc | 0.773 |
| Pharm assessed by | IChem |

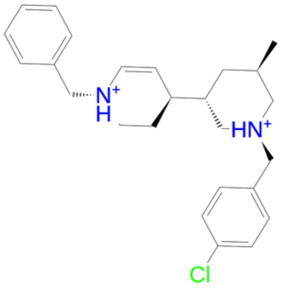

| | |
|---|---|
| Name | FJRL-94 |
| SIFt Tc | 0.773 |
| Pharm assessed by | IChem |

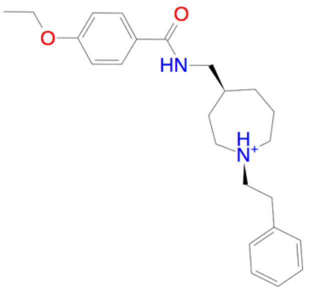

| | |
|---|---|
| Name | FJRL-95 |
| SIFt Tc | 0.773 |
| Pharm assessed by | IChem |

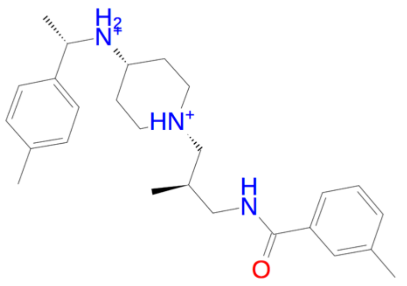

| | |
|---|---|
| Name | FJRL-96 |
| SIFt Tc | 0.773 |
| Pharm assessed by | IChem |

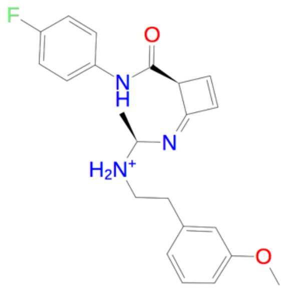

| | |
|---|---|
| Name | FJRL-97 |
| SIFt Tc | 0.762 |
| Pharm assessed by | IChem |

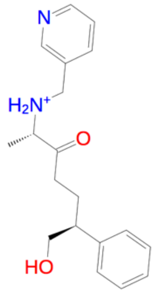

| | |
|---|---|
| Name | FJRL-98 |
| SIFt Tc | 0.762 |
| Pharm assessed by | USRCAT |

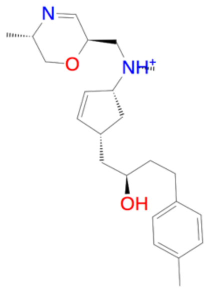

| | |
|---|---|
| Name | FJRL-99 |
| SIFt Tc | 0.762 |
| Pharm assessed by | IChem |

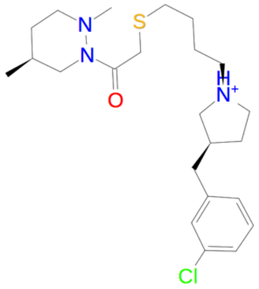

| | |
|---|---|
| Name | FJRL-100 |
| SIFt Tc | 0.762 |
| Pharm assessed by | IChem |

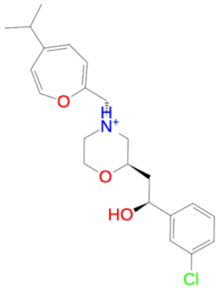

| | |
|---|---|
| Name | FJRL-101 |
| SIFt Tc | 0.762 |
| Pharm assessed by | IChem |

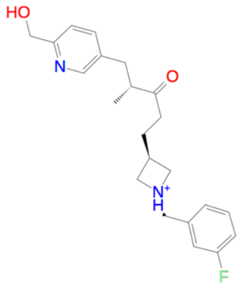

| | |
|---|---|
| Name | FJRL-102 |
| SIFt Tc | 0.762 |
| Pharm assessed by | USRCAT |

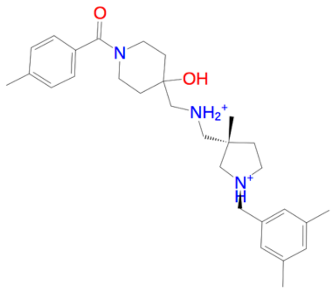

| | |
|---|---|
| Name | FJRL-103 |
| SIFt Tc | 0.762 |
| Pharm assessed by | IChem |

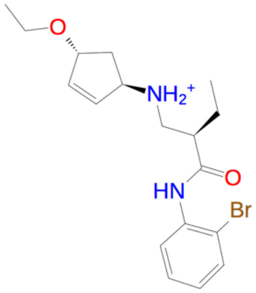

| | |
|---|---|
| Name | FJRL-104 |
| SIFt Tc | 0.762 |
| Pharm assessed by | IChem |

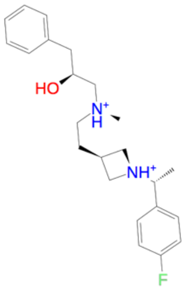

| | |
|---|---|
| Name | FJRL-105 |
| SIFt Tc | 0.762 |
| Pharm assessed by | USRCAT |

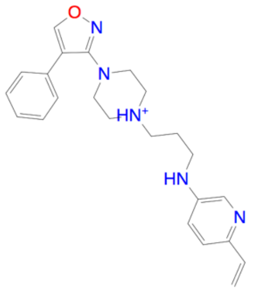

| | |
|---|---|
| Name | FJRL-106 |
| SIFt Tc | 0.762 |
| Pharm assessed by | IChem |

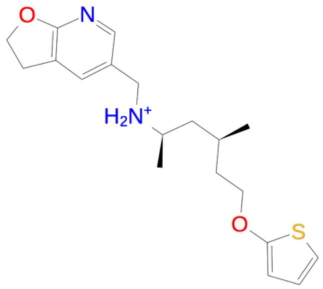

| | |
|---|---|
| Name | FJRL-107 |
| SIFt Tc | 0.762 |
| Pharm assessed by | IChem |

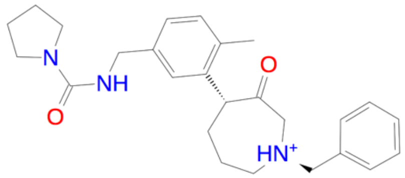

| | |
|---|---|
| Name | FJRL-108 |
| SIFt Tc | 0.762 |
| Pharm assessed by | IChem |

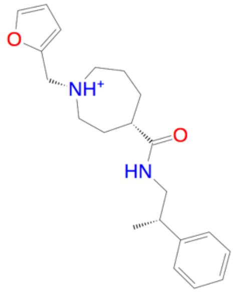

| | |
|---|---|
| Name | FJRL-109 |
| SIFt Tc | 0.762 |
| Pharm assessed by | USRCAT |

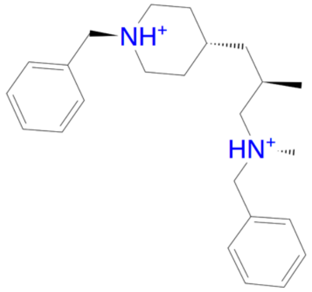

| | |
|---|---|
| Name | FJRL-110 |
| SIFt Tc | 0.762 |
| Pharm assessed by | IChem |

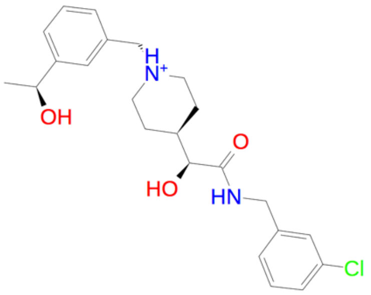

| | |
|---|---|
| Name | FJRL-111 |
| SIFt Tc | 0.762 |
| Pharm assessed by | USRCAT |

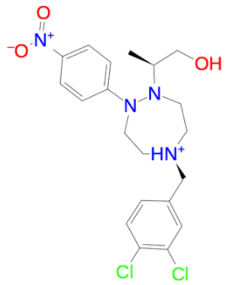

| | |
|---|---|
| Name | FJRL-112 |
| SIFt Tc | 0.762 |
| Pharm assessed by | IChem |

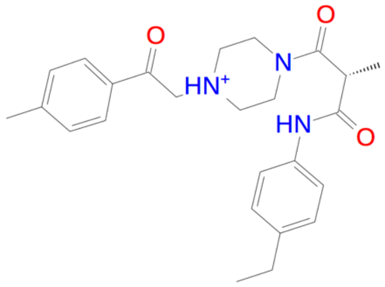

| | |
|---|---|
| Name | FJRL-113 |
| SIFt Tc | 0.762 |
| Pharm assessed by | IChem |

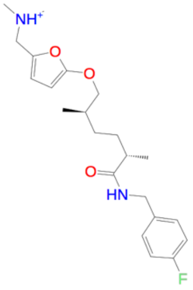

| | |
|---|---|
| Name | FJRL-114 |
| SIFt Tc | 0.762 |
| Pharm assessed by | USRCAT |

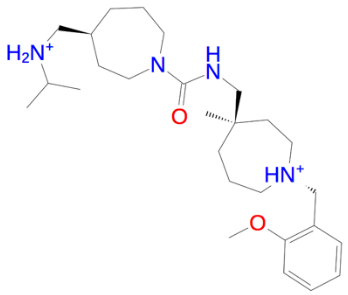

| | |
|---|---|
| Name | FJRL-115 |
| SIFt Tc | 0.762 |
| Pharm assessed by | IChem |

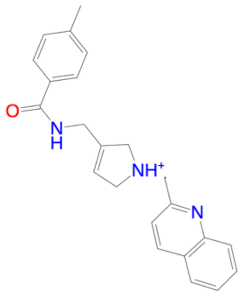

| | |
|---|---|
| Name | FJRL-116 |
| SIFt Tc | 0.762 |
| Pharm assessed by | IChem |

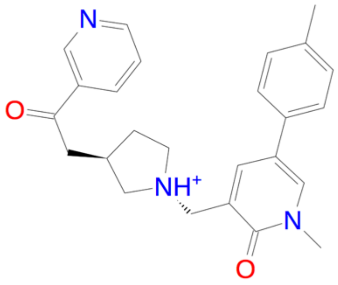

| | |
|---|---|
| Name | FJRL-117 |
| SIFt Tc | 0.762 |
| Pharm assessed by | IChem |

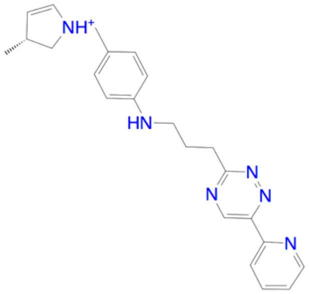

| | |
|---|---|
| Name | FJRL-118 |
| SIFt Tc | 0.762 |
| Pharm assessed by | IChem |

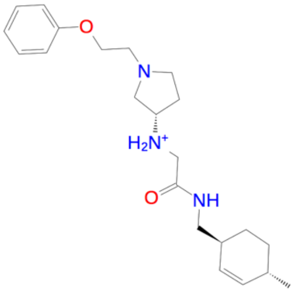

| | |
|---|---|
| Name | FJRL-119 |
| SIFt Tc | 0.762 |
| Pharm assessed by | IChem |

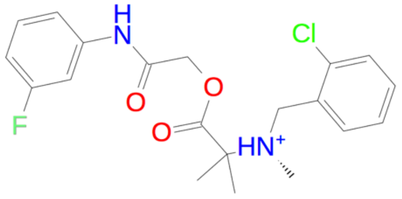

| | |
|---|---|
| Name | FJRL-120 |
| SIFt Tc | 0.762 |
| Pharm assessed by | USRCAT |

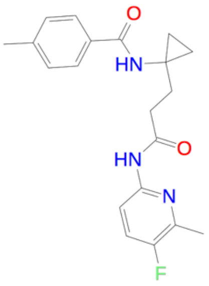

| | |
|---|---|
| Name | FJRL-121 |
| SIFt Tc | 0.762 |
| Pharm assessed by | USRCAT |

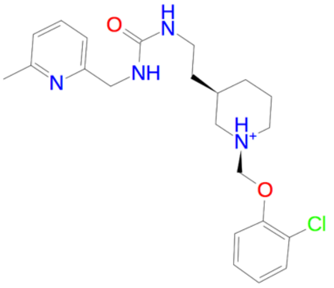

| | |
|---|---|
| Name | FJRL-122 |
| SIFt Tc | 0.762 |
| Pharm assessed by | IChem |

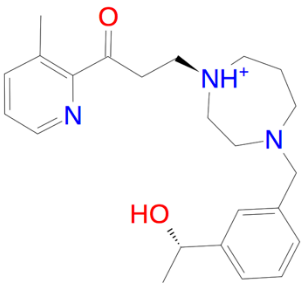

| | |
|---|---|
| Name | FJRL-123 |
| SIFt Tc | 0.762 |
| Pharm assessed by | USRCAT |

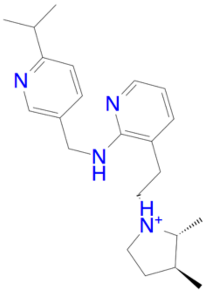

| | |
|---|---|
| Name | FJRL-124 |
| SIFt Tc | 0.762 |
| Pharm assessed by | IChem |

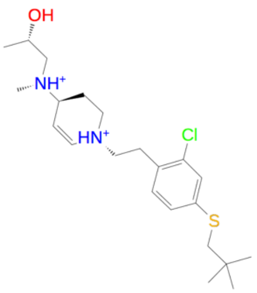

| | |
|---|---|
| Name | FJRL-125 |
| SIFt Tc | 0.762 |
| Pharm assessed by | IChem |

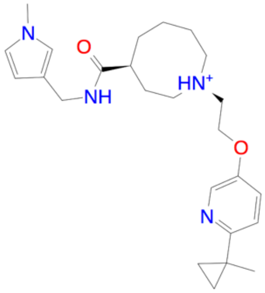

| | |
|---|---|
| Name | FJRL-126 |
| SIFt Tc | 0.762 |
| Pharm assessed by | IChem |

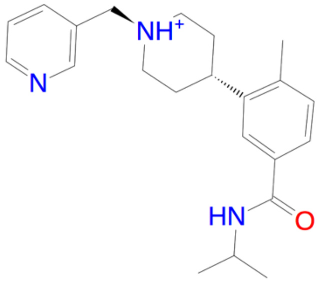

| | |
|---|---|
| Name | FJRL-127 |
| SIFt Tc | 0.762 |
| Pharm assessed by | IChem |

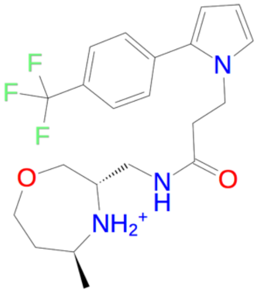

| | |
|---|---|
| Name | FJRL-128 |
| SIFt Tc | 0.762 |
| Pharm assessed by | IChem |

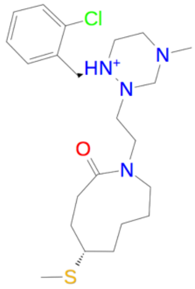

| | |
|---|---|
| Name | FJRL-129 |
| SIFt Tc | 0.762 |
| Pharm assessed by | IChem |

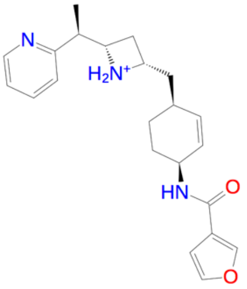

| | |
|---|---|
| Name | FJRL-130 |
| SIFt Tc | 0.762 |
| Pharm assessed by | IChem |

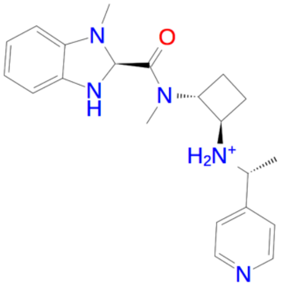

| | |
|---|---|
| Name | FJRL-131 |
| SIFt Tc | 0.762 |
| Pharm assessed by | USRCAT |

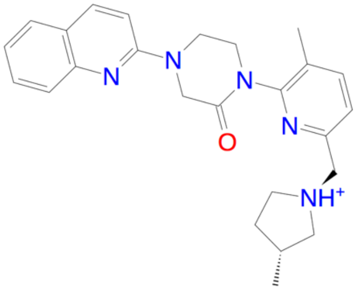

| | |
|---|---|
| Name | FJRL-132 |
| SIFt Tc | 0.762 |
| Pharm assessed by | IChem |

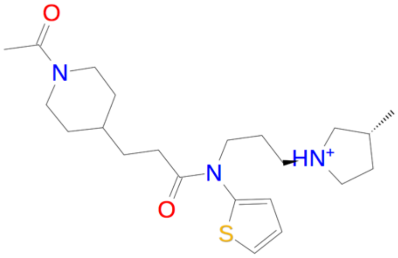

| | |
|---|---|
| Name | FJRL-133 |
| SIFt Tc | 0.762 |
| Pharm assessed by | IChem |

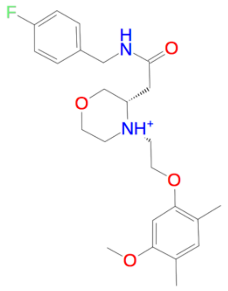

| | |
|---|---|
| Name | FJRL-134 |
| SIFt Tc | 0.762 |
| Pharm assessed by | USRCAT |

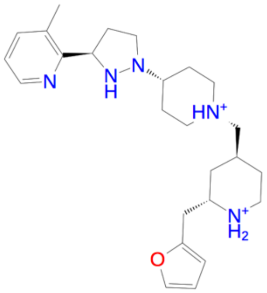

| | |
|---|---|
| Name | FJRL-135 |
| SIFt Tc | 0.762 |
| Pharm assessed by | IChem |

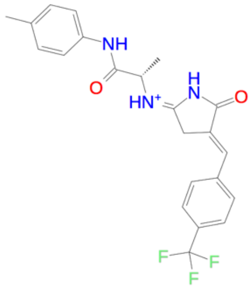

| | |
|---|---|
| Name | FJRL-136 |
| SIFt Tc | 0.762 |
| Pharm assessed by | IChem |

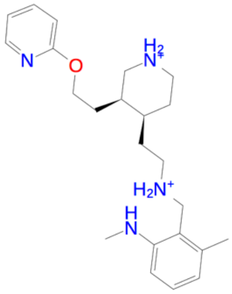

| | |
|---|---|
| Name | FJRL-137 |
| SIFt Tc | 0.762 |
| Pharm assessed by | IChem |

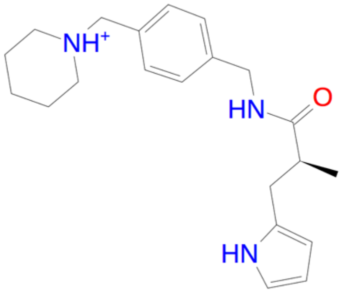

| | |
|---|---|
| Name | FJRL-138 |
| SIFt Tc | 0.762 |
| Pharm assessed by | USRCAT |

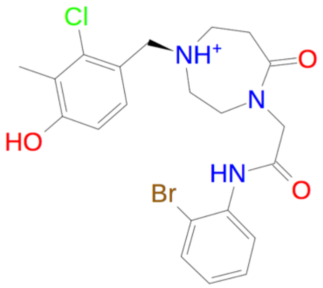

| | |
|---|---|
| Name | FJRL-139 |
| SIFt Tc | 0.762 |
| Pharm assessed by | USRCAT |

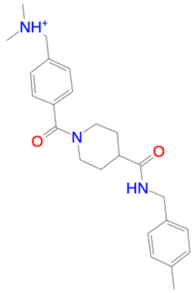

| | |
|---|---|
| Name | FJRL-140 |
| SIFt Tc | 0.762 |
| Pharm assessed by | IChem |

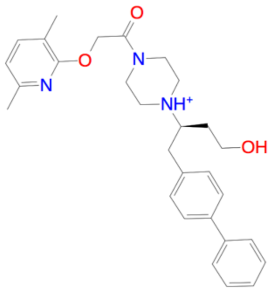

| | |
|---|---|
| Name | FJRL-141 |
| SIFt Tc | 0.76 |
| Pharm assessed by | IChem |

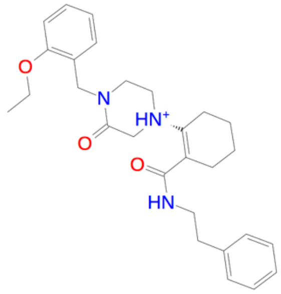

| | |
|---|---|
| Name | FJRL-142 |
| SIFt Tc | 0.75 |
| Pharm assessed by | IChem |

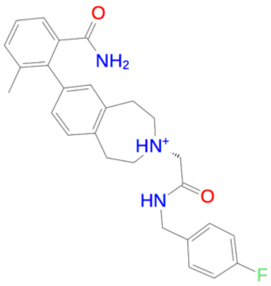

| | |
|---|---|
| Name | FJRL-143 |
| SIFt Tc | 0.75 |
| Pharm assessed by | IChem |

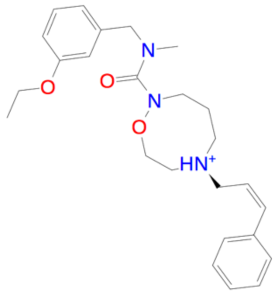

| | |
|---|---|
| Name | FJRL-144 |
| SIFt Tc | 0.75 |
| Pharm assessed by | IChem |

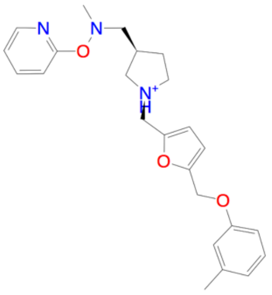

| | |
|---|---|
| Name | FJRL-145 |
| SIFt Tc | 0.75 |
| Pharm assessed by | IChem |

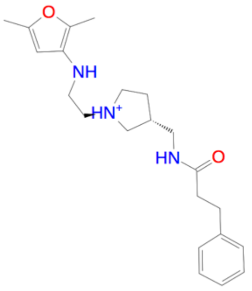

| | |
|---|---|
| Name | FJRL-146 |
| SIFt Tc | 0.75 |
| Pharm assessed by | USRCAT |

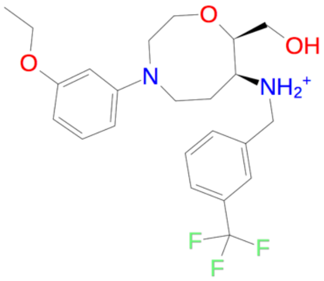

| | |
|---|---|
| Name | FJRL-147 |
| SIFt Tc | 0.75 |
| Pharm assessed by | IChem |

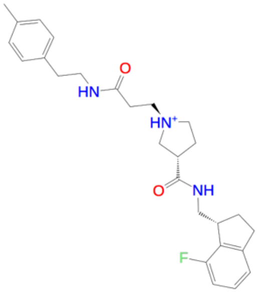

| | |
|---|---|
| Name | FJRL-148 |
| SIFt Tc | 0.75 |
| Pharm assessed by | IChem |

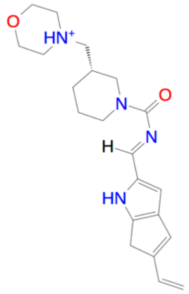

| | |
|---|---|
| Name | FJRL-149 |
| SIFt Tc | 0.75 |
| Pharm assessed by | IChem |

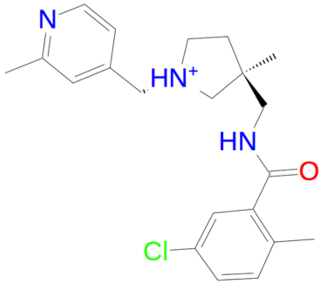

| | |
|---|---|
| Name | FJRL-150 |
| SIFt Tc | 0.75 |
| Pharm assessed by | USRCAT |

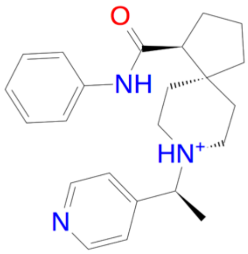

| | |
|---|---|
| Name | FJRL-151 |
| SIFt Tc | 0.75 |
| Pharm assessed by | USRCAT |

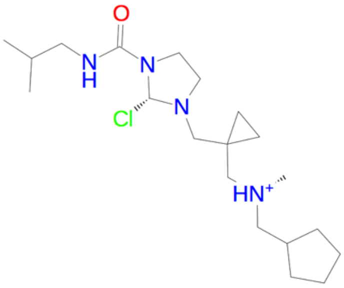

| | |
|---|---|
| Name | FJRL-152 |
| SIFt Tc | 0.75 |
| Pharm assessed by | IChem |

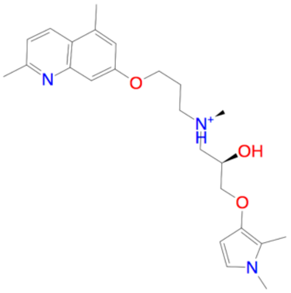

| | |
|---|---|
| Name | FJRL-153 |
| SIFt Tc | 0.75 |
| Pharm assessed by | IChem |

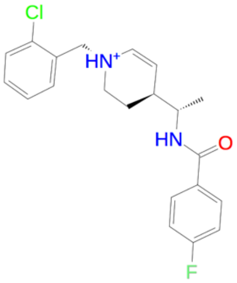

| | |
|---|---|
| Name | FJRL-154 |
| SIFt Tc | 0.75 |
| Pharm assessed by | USRCAT |

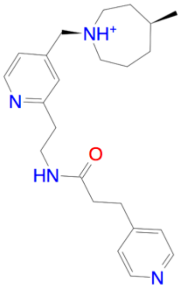

| | |
|---|---|
| Name | FJRL-155 |
| SIFt Tc | 0.75 |
| Pharm assessed by | USRCAT |

| | | | | | |
|---|---|---|---|---|---|
| 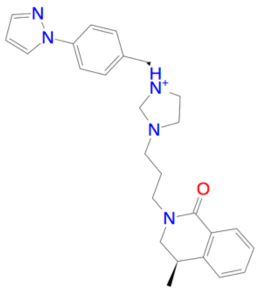 | | 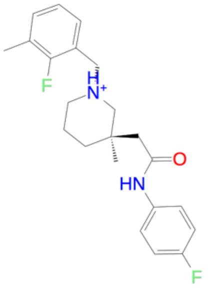 | | 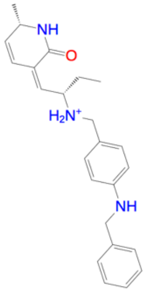 | |
| Name | FJRL-156 | Name | FJRL-157 | Name | FJRL-158 |
| SIFt Tc | 0.75 | SIFt Tc | 0.75 | SIFt Tc | 0.75 |
| Pharm assessed by | IChem | Pharm assessed by | USRCAT | Pharm assessed by | IChem |
| 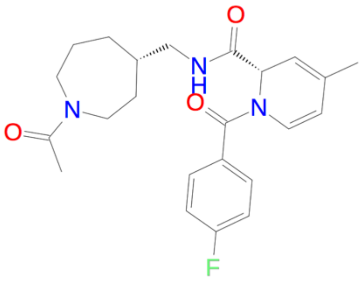 | | 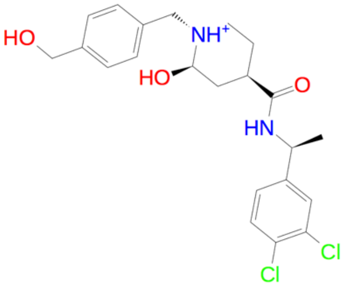 | | 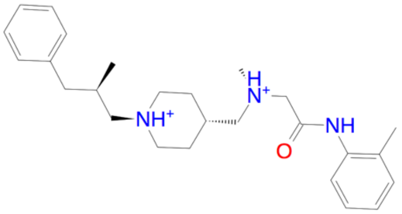 | |
| Name | FJRL-159 | Name | FJRL-160 | Name | FJRL-161 |
| SIFt Tc | 0.75 | SIFt Tc | 0.75 | SIFt Tc | 0.75 |
| Pharm assessed by | IChem | Pharm assessed by | USRCAT | Pharm assessed by | IChem |
| 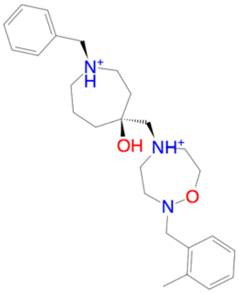 | | 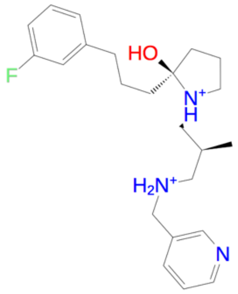 | | 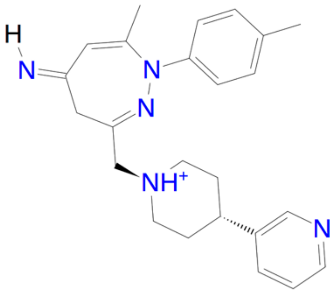 | |
| Name | FJRL-162 | Name | FJRL-163 | Name | FJRL-164 |
| SIFt Tc | 0.75 | SIFt Tc | 0.75 | SIFt Tc | 0.75 |
| Pharm assessed by | IChem | Pharm assessed by | USRCAT | Pharm assessed by | USRCAT |
| 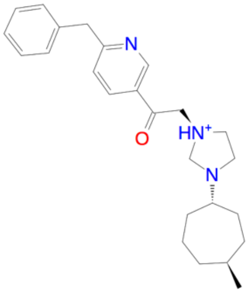 | | 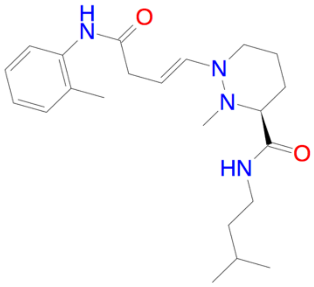 | | 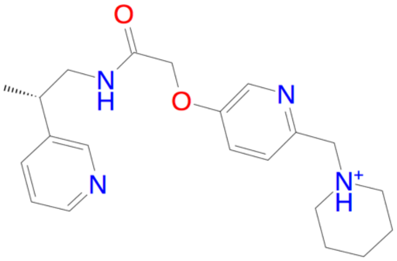 | |
| Name | FJRL-165 | Name | FJRL-166 | Name | FJRL-167 |
| SIFt Tc | 0.75 | SIFt Tc | 0.75 | SIFt Tc | 0.75 |
| Pharm assessed by | IChem | Pharm assessed by | IChem | Pharm assessed by | IChem |

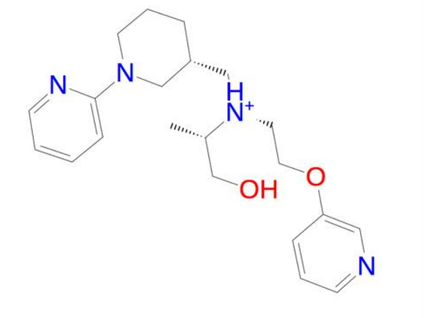

| | |
|---|---|
| Name | FJRL-168 |
| SIFt Tc | 0.75 |
| Pharm assessed by | USRCAT |

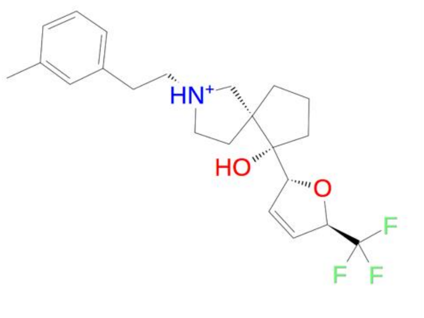

| | |
|---|---|
| Name | FJRL-169 |
| SIFt Tc | 0.75 |
| Pharm assessed by | IChem |

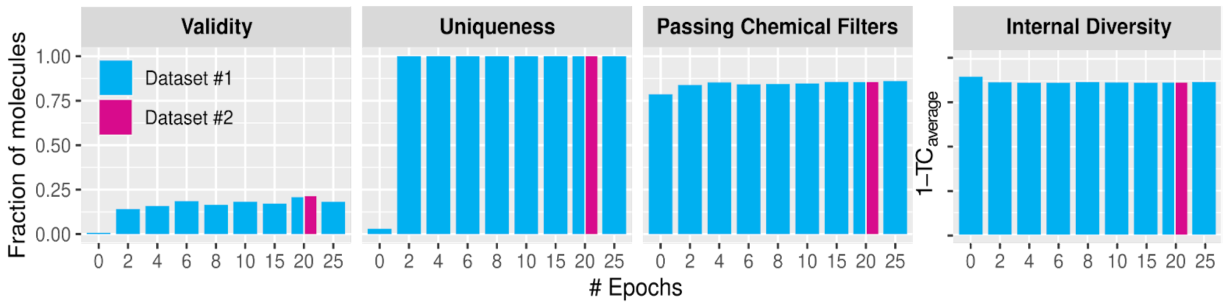


**Figure S1. Performance of the deep generative model in producing chemically valid, novel, and diverse molecules during pre-training.** Validity: fraction of valid SMILES as assessed by their successful conversion to RDKit and OpenEye molecular objects; Uniqueness: fraction of molecules that are not duplicated; Passing Chemical Filters: fraction of molecules that only have C, N, S, O, F, Cl, Br, and H atoms, cycles with less than 8 atoms, and pass both PAINS filters and MCFs; Internal Diversity: chemical diversity as per 1 - $Tc_{average}$ calculated using Morgan fingerprints. An epoch consists of one pass over the entire Dataset #1.

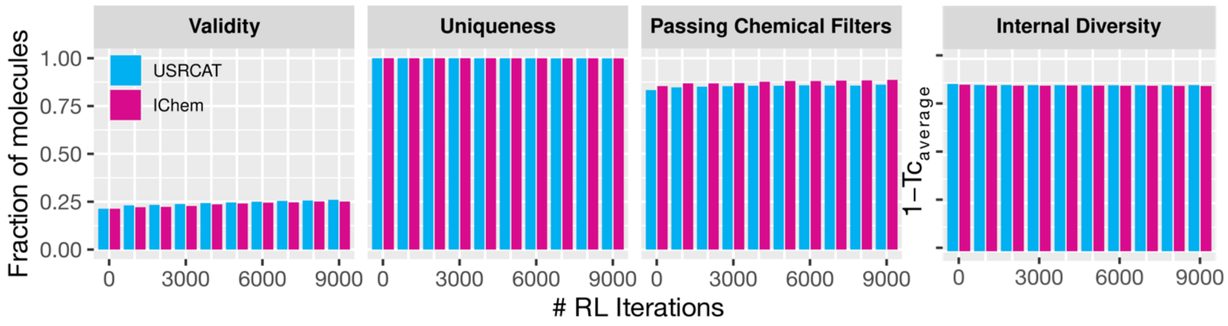


**Figure S2. Performance of the deep generative model in producing chemically valid, novel, and diverse molecules during reinforcement learning.** Validity: fraction of valid SMILES as assessed by their successful conversion to RDKit and OpenEye molecular objects; Uniqueness: fraction of molecules that are not duplicated; Passing Chemical Filters: fraction of molecules that only have C, N, S, O, F, Cl, Br, and H atoms, cycles with less than 8 atoms, and pass both PAINS filters and MCFs; Internal Diversity: chemical diversity as per 1 - $Tc_{average}$ calculated using Morgan fingerprints. An iteration consists of generation and scoring one batch of 200 SMILES.

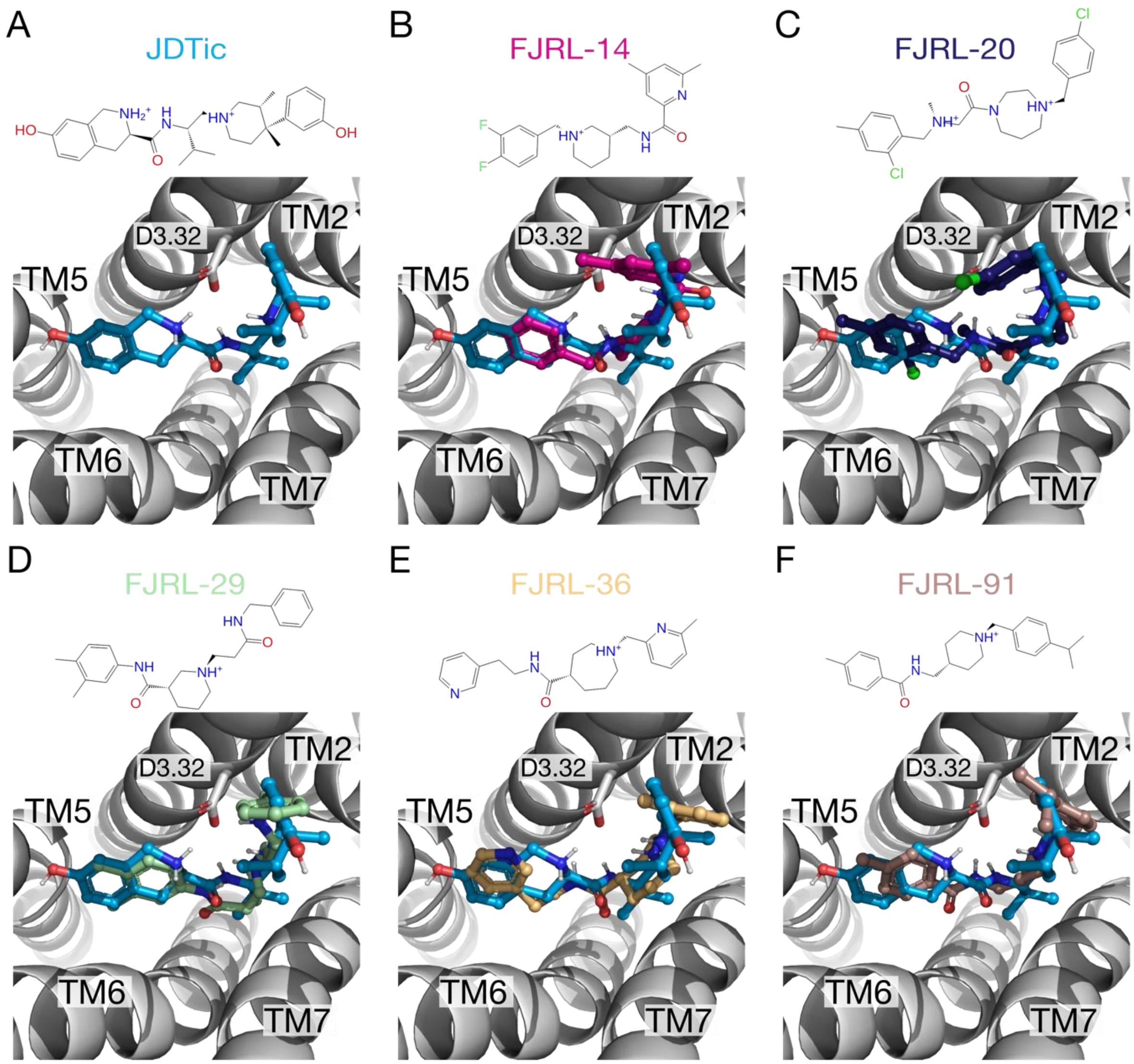


**Figure S3. Compounds prioritized for chemical synthesis docked at KOR and compared to JDTic's binding mode.** A-F) KOR is depicted in a cartoon representation and colored white, with residue D3.32 shown in sticks. JDTic and the prioritized molecules are also shown in ball-and-stick representation and colored cyan (JDTic), magenta (FJRL-14), deep violet (FJRL-20), pale green (FJRL-29), pale orange (FJRL-36), and dark salmon (FJRL-91), respectively.

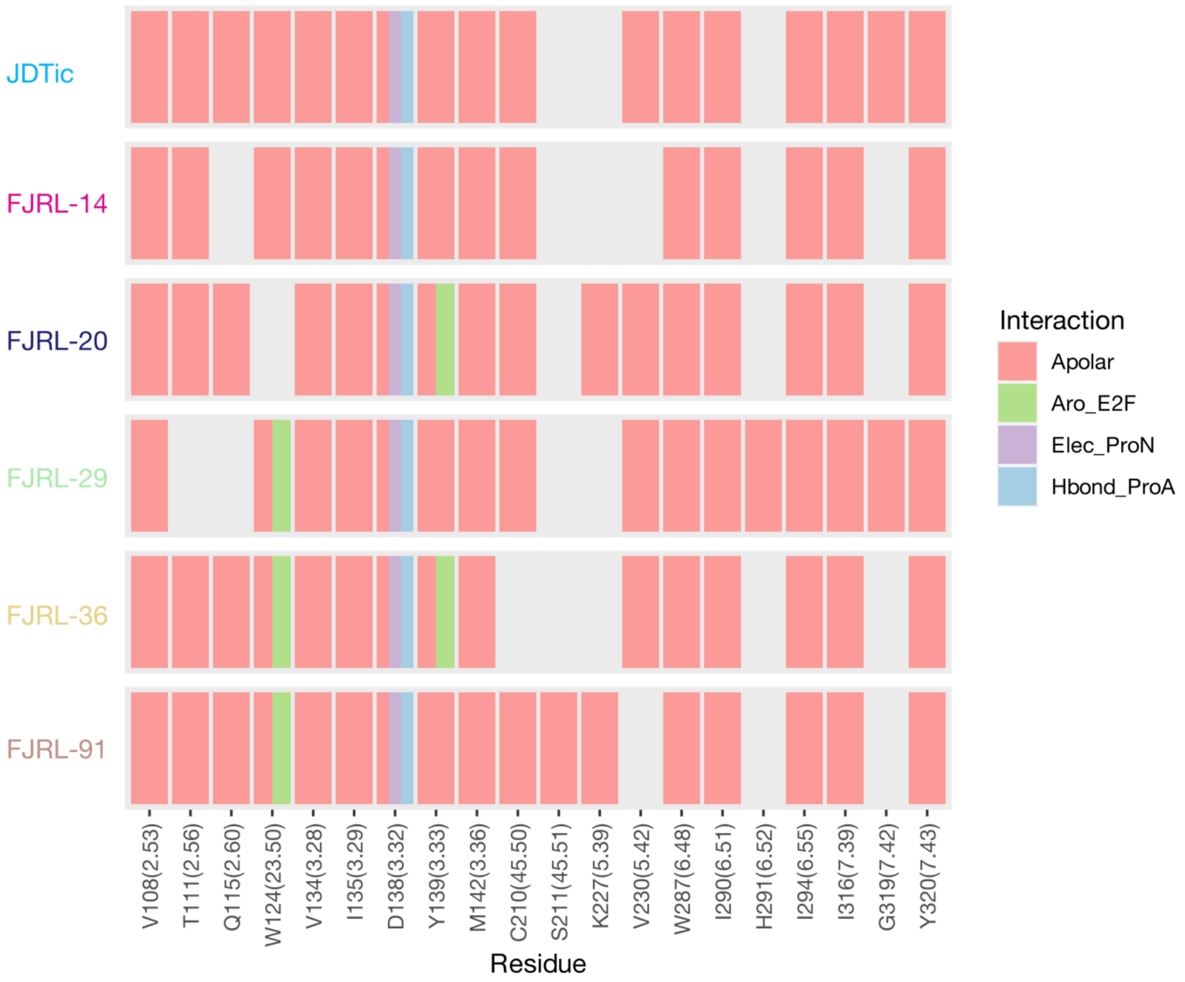


**Figure S4. Protein-ligand interactions of compounds prioritized for chemical synthesis docked at KOR and compared to JDTic's.** Structural interactions fingerprints of JDTic and the prioritized molecules at KOR. Apolar: carbon-carbon interactions; Aro_E2F: face-to-face aromatic interactions; Elec_ProN: electrostatic interaction with the protein residue negatively charged; and Hbond_ProA: hydrogen-bonds with a protein residue as the acceptor. The Ballesteros-Weinstein generic numbering[43] of each residue is indicated in parenthesis.
